# Long-read and chromosome-scale assembly of the hexaploid wheat genome achieves high resolution for research and breeding

**DOI:** 10.1101/2021.08.24.457458

**Authors:** Jean-Marc Aury, Stefan Engelen, Benjamin Istace, Cécile Monat, Pauline Lasserre-Zuber, Caroline Belser, Corinne Cruaud, Hélène Rimbert, Philippe Leroy, Sandrine Arribat, Isabelle Dufau, Arnaud Bellec, David Grimbichler, Nathan Papon, Etienne Paux, Marion Ranoux, Adriana Alberti, Patrick Wincker, Frédéric Choulet

**Affiliations:** Génomique Métabolique, Genoscope, Institut François Jacob, CEA, CNRS, Univ Evry, Université Paris-Saclay, 91057 Evry, France; GDEC, Université Clermont Auvergne, INRAE, UMR1095, 63000 Clermont-Ferrand, France; Commissariat à l’Energie Atomique (CEA), Institut François Jacob, Genoscope, F-91057 Evry, France; INRAE, CNRGV French Plant Genomic Resource Center, F-31320, Castanet Tolosan, France; Mésocentre Clermont Auvergne, DOSI / Bâtiment Turing, 7 avenue Blaise Pascal, 63178 Aubière CEDEX; Université Paris-Saclay, CEA, CNRS, Institute for Integrative Biology of the Cell (I2BC), 91198, Gif-sur-Yvette, France

## Abstract

The sequencing of the wheat (*Triticum aestivum*) genome has been a methodological challenge for many years due to its large size (15.5 Gb), repeat content, and hexaploidy. Many initiatives aiming at obtaining a reference genome of cultivar Chinese Spring have been launched in the past years and it was achieved in 2018 as the result of a huge effort to combine short-read sequencing with many other resources. Reference-quality genome assemblies were then produced for other accessions but the rapid evolution of sequencing technologies offers opportunities to reach high-quality standards at lower cost. Here, we report on an optimized procedure based on long-reads produced on the ONT (Oxford Nanopore Technology) PromethION device to assemble the genome of the French bread wheat cultivar Renan. We provide the most contiguous and complete chromosome-scale assembly of a bread wheat genome to date. Coupled with an annotation based on RNA-Seq data, this resource will be valuable for the crop community and will facilitate the rapid selection of agronomically important traits. We also provide a framework to generate high-quality assemblies of complex genomes using ONT.

## Introduction

Bread wheat (*Triticum aestivum*) is among the most important cereal crops and a better knowledge in the area of wheat genomics is needed to face the main challenge of ensuring food security to a growing population in the context of climate change. Improving productivity requires both that local producers adapt their practices to increase their climate resilience and a better understanding of the wheat production systems. In this context, a better knowledge of the wheat genome and its gene content, but also the sequencing of numerous accessions, are essential.

However, the genome of bread wheat is particularly characterized by its complexity. Indeed, this hexaploid genome is the result of two interspecific hybridization events. The earliest cultivated wheat was diploid, but humans have intensified the cultivation of polyploid species. Recent studies show that these polyploid species appear to be advantaged by their genomic plasticity[1]. Indeed, modifications of the gene space and related elements are buffered by the polyploid nature of wheat and open a wider field to selection. Bread wheat is composed of three subgenomes A, B and D derived from three ancestral diploid species that diverged between 2.5 and 6 million years ago[2].

The wheat genome is one of the largest among sequenced plant genomes (15.5 Gb), mainly composed of repetitive sequences (ca. >85%), and contains many homoeologous regions between the three subgenomes (A, B and D). Repetitive sequences and polyploidy pose serious challenges in the generation of genome assemblies. The adventure of sequencing the hexaploid wheat genome began in 2005 with the creation of the International Wheat Genome Sequencing Consortium (IWGSC)[3]. With the advent of sequencing technologies, the wheat genome has been competitively sequenced several times[4–6]. The first reference-quality genome sequence with a comprehensive annotation was published by the IWGSC in August 2018[7] for the accession Chinese Spring (CS). This assembly represents a tremendous resource for the scientific community and offers the promise of facilitating and accelerating breeding efforts.

More recently, fifteen genomes of hexaploid wheat have been published[8] which represents a new step in the knowledge of the wheat model. Ten of these new wheat genomes have been assembled at the chromosome level, allowing for comparative analysis on a scale that was previously impossible. While being a valuable and highly validated resource using multiple technologies, these assemblies were produced using short-read technologies and therefore may contain a higher number of gaps compared to genomes assembled with long reads[9–13]. In 2017, an assembly of the CS genome using long-reads was produced[5], although not annotated, highlighting the added-value of long-reads in such complex genomes. By accumulating long-read assemblies, the scientific community is now aware of the flaw in short-read strategies. Indeed they underestimate the repetitive content of the genome and more importantly can lack tandemly duplicated genes[14,15]. Several years ago, Pacific Biosciences (PACBIO) and Oxford Nanopore (ONT) sequencing technologies were commercialized with the promise to sequence long DNA fragments and revolutionize complex genome assemblies.

Here, we report the first hexaploid wheat genome based on ONT long-reads. We sequenced the genome of the French variety Renan, one of the most used varieties in organic farming. The Renan genome carries multiple resistance genes against fungal pathogens (leaf rust, stem rust, yellow rust, eyespot) originating from introgression of DNA regions coming from the wild species *Aegilops ventricosa*. We used the PromethION device and organized the assembled contigs at the chromosome scale using optical maps (BioNano Genomics, BNG) and Hi-C libraries (Arima Genomics, AG). This assembly has a contig N50 of 2.2 Mb, which is a 30-fold improvement over existing chromosome-scale assemblies.

## Results

### Genome sequencing and optical maps

We sequenced genomic DNA using 20 ONT flow cells (2 MinION and 18 PromethION) which produced 12M reads representing 1.1 Tb. All the reads were originally base called using the guppy 2.0 software, but given the improvement of guppy software during our project, we decided to call bases using a newer version of the guppy software (version 3.6 with High Accuracy setting). This dataset represented a coverage of 63x of the hexaploid wheat genome and the read N50 was of 24.6kb. More importantly, we got 3.1M reads larger than 50kb representing a 14x genome coverage (Table S1). In addition, we generated Illumina short-reads and long-range data for respectively polishing and organizing nanopore contigs. We produced an optical map using the Saphyr instrument commercialized by Bionano Genomics (BNG). High molecular weight DNA was extracted and labeled using the Direct Label and Stain Chemistry (DLS) with the DLE-1 enzyme. The DLE-1 optical map was assembled using proprietary tools provided by BNG and had a cumulative size of 14.9 Gb with an N50 of 37.5 Mb (Table S2). Four Hi-C libraries from two biological replicates were prepared using the Arima Genomics protocol and sequenced on an Illumina sequencer to reach 537 Gb i.e., a depth of 35x. We used a sample of 240 million read pairs (72 Gb, 5x) to build a Hi-C map.

### Genome assembly

Since the dataset was too large for many long-read assemblers, we sampled a 30x coverage by selecting the longest reads (Table S1). This subset was assembled using multiple assembly tools dedicated to processing this large amount of data (Redbean[16], SMARTdenovo[17] and Flye[18]). SMARTdenovo is not among the fastest algorithms and has not been updated for several years, but since it can be easily parallelized, it remains an interesting choice for assembling large genomes. The overlap and consensus calculations were split into 60 chunks and each were run on a 32-core server and took about two days and ten hours respectively. In comparison, Redbean was able to generate an assembly after just seven days on a 64-core server with 3TB of memory while Flye needed 43 days on the same computer server. Surprisingly, the redbean assembly had a cumulative size two times higher than the expected genome size (29.6Gb vs 14.5Gb), a low contiguity and contained a large amount of short contigs. The SMARTdenovo and Flye assemblies were highly comparable, but Flye was the most contiguous (contigs N50 of 1.8 Mb vs 1.1 Mb) and SMARTdenovo had a cumulative size closer to the expected one (14.1 Gb vs 13.0 Gb, Table S3). Additionally, even though the assemblies were polished later, the raw SMARTdenovo assembly contained a higher number of complete BUSCO genes (83.0% vs 49.5%) which indicates that its consensus module is more efficient.

The SMARTdenovo and Flye assemblies were successively polished using Racon[19] and Medaka (https://github.com/nanoporetech/medaka) with long reads and Hapo-G[20] with short reads. Polished contigs were validated and organized into scaffolds using the DLE-1 optical map and proprietary tools provided by BNG. As expected, due to its lower cumulative size, Flye scaffolds contained a larger proportion of unknown bases (851 Mb and 262 Mb). Based on these results (proportion of gaps and gene completion), the assembly produced by SMARTdenovo[17] was selected (Table S4). Local contig duplications (negative gaps) were resolved using BiSCoT^22^, which improved the contigs N50 from 1.2 Mb up to 2.1 Mb. Finally, the resulting assembly was polished one last time using Hapo-G[20] with short reads. This led to 2,904 scaffolds (larger than 30kb) representing 14.26 Gb with a N50 of 48 Mb (79 scaffolds) and a maximum scaffold size of 254 Mb. Thus, the genome size is in the same range as all other available reference quality assemblies of *T. aestivum:* e.g. 14.29 Gb for *cv*. LongReach Lancer, 14.55 Gb for *cv*. Chinese Spring, and 14.96 Gb for *cv*. SY Mattis.

### Construction and validation of pseudomolecules

We then guided the construction of the 21 chromosome sequences (i.e. pseudomolecules) based on collinearity with the CS (Chinese Spring) RefSeq Assembly v2.1[22]. Given the complexity of this hexaploid genome, we established a dedicated approach in order to anchor each Renan scaffold based on similarity search against CS. To avoid problems due to multiple mappings, we selected a dataset of uniquely mappable sequences. Genes are not uniquely mappable since most of them are repeated as three homoeologous copies sharing on average 97% nucleotide identity. In addition, the gene density (1 gene every 130kb on average) is too low to anchor small Renan scaffolds that do not carry genes. Thus, we used 150 bp tags corresponding to the 5’ and 3’ junctions between a transposable element (TE) and its insertion site (75 bps on each side) which are called ISBP (Insertion Site-Based Polymorphism) markers and are highly abundant and uniquely mappable in the wheat genome[23]. We designed a dataset of 5.76 million ISBPs from CS assembly which represent 1 ISBP every 2.5kb. Their mapping enabled the anchoring of 2,566 scaffolds on 21 pseudomolecules representing 14.20 Gb (99% of the assembly). We then used Hi-C data to validate the assembly and to correct the mis-ordered and mis-oriented scaffolds. The Hi-C map revealed only a few inconsistencies, demonstrating that the collinearity between CS and Renan was strong enough to guide the anchoring in a very accurate manner. The Hi-C map-based curation led to the detection of 18 chimeric scaffolds that were split into 2 or 3 pieces and to the correction of the location and/or orientation of 198 scaffolds. The final assembly was composed of 21 pseudomolecules (Figure 1) with 338 unanchored scaffolds representing 61 Mb only.

**Figure 1.**
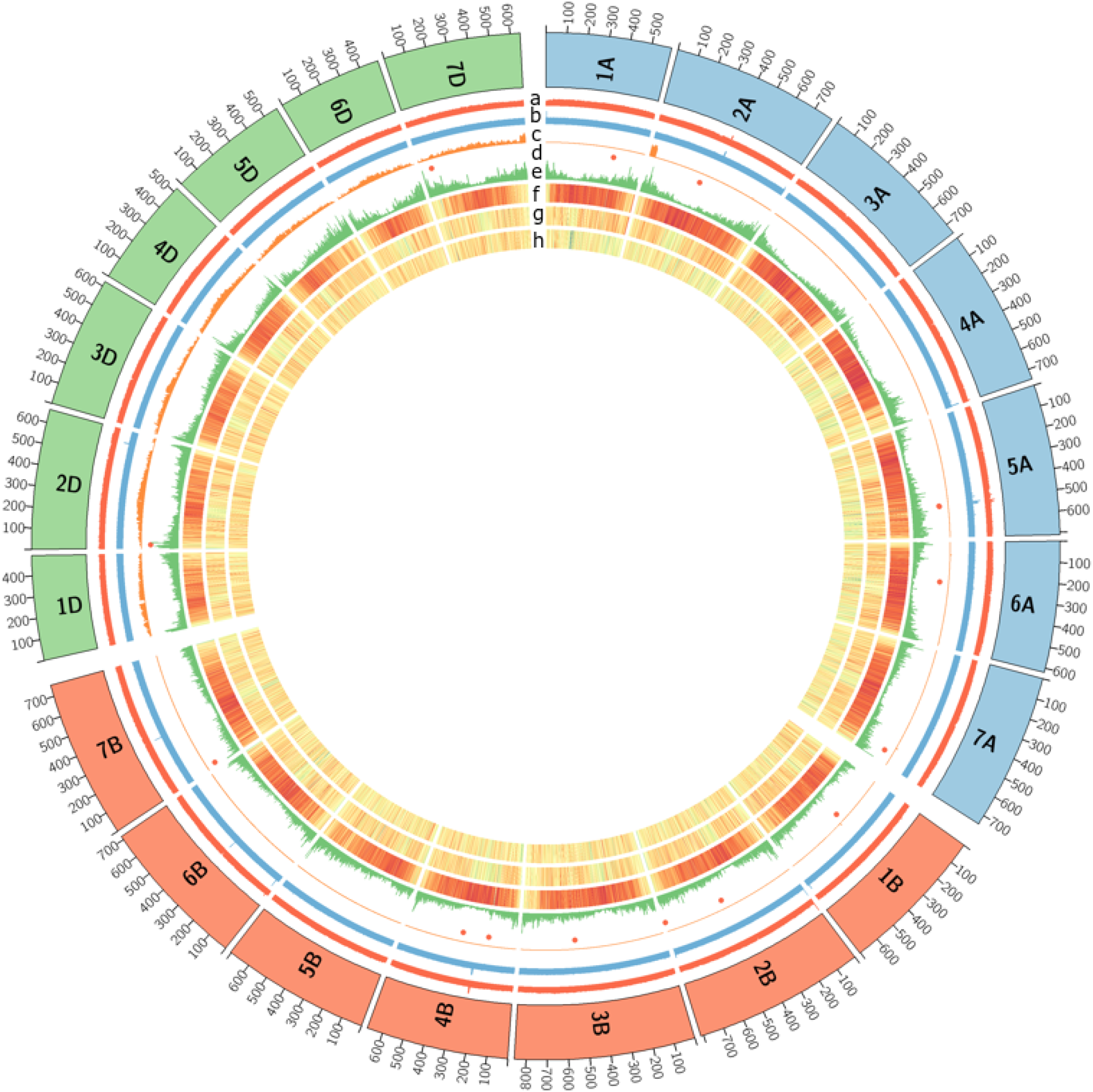
Genome overview of the 21 chromosomes of hexaploid *T. aestivum* Renan (the 7 A chromosomes are in blue, the 7 B chromosomes in orange and the 7 D chromosomes in green). From inner to outer track: (a) Coverage with short reads, (b) Coverage with long reads, (c) coverage with *Ae. ventricosa* short reads, (d) Red dots represent large deletions (>500Kb), (e) Gene density, (f) Density of CACTA (DNA transposon) elements, (g) Density of Copia elements, (h) Density of Gypsy elements. All densities and coverage are calculated in 1-Mb windows; yellow and red colors in density plots indicate lower and higher values, respectively.

### Quality assessment of the assembly

First, we calculated the overall quality of the sequence using Merqury and Illumina reads. We obtained an average quality value (QV) of 32.8, a lower QV than that obtained with short-reads assemblies, but consistent with QV already reported for plant genomes sequenced by ONT[24]. Indeed, using Illumina reads and the CS RefSeq v2.1 assembly, Merqury computed a QV of 44.5 (Table 1). This shows that per-base quality is still an issue, at least with the version of the technology used in this study. However, this could be tempered by the fact that coding regions, due to lower repeating regions, may have higher precision.

**Table 1:** Comparison of *Triticum aestivum L*. genome assemblies. *NG50 and NG90 were calculated using a genome size of 15Gb.

|  |  | Renan<br>This study | Chinese Spring<br>RefSeq_v2.1<br>from Zhu et al.[22] |
| --- | --- | --- | --- |
| Number of contigs |  | 12,982 | 306,746 |
| Cumulative size (bp) |  | 14,001,122,256 | 14,317,423,665 |
| N50 (bp) |  | 2,159,703 | 341,062 |
| L50 |  | 1,958 | 12,223 |
| N90 (bp) |  | 598,285 | 32,302 |
| L90 |  | 6,645 | 59,261 |
| NG50* (bp) |  | 1,973,000 | 322,161 |
| LG50 |  | 2,202 | 13,254 |
| NG90* (bp) |  | 264,272 | 16,550 |
| LG90 |  | 8,816 | 85,688 |
| Longest contig (bp) |  | 15,116,687 | 3,528,546 |
| Number of chromosomes |  | 21 | 21 |
| Cumulative size (bp) |  | 14,195,643,615 | 14,225,829,371 |
| N50 (bp) |  | 703,299,328 | 713,360,512 |
| L50 |  | 10 | 10 |
| N90 (bp) |  | 520,815,552 | 518,332,608 |
| L90 |  | 19 | 19 |
| Longest (bp) |  | 854,463,248 | 851,934,019 |
| % of N |  | 1.78% | 1.52% |
| BUSCO on assemblies<br>(N=4,896) | Complete | 99.1% | 99.3% |
|  | Duplicated | 94.7% | 96.1% |
|  | Fragmented | 0.1% | 0.1% |
|  | Missing | 0.8% | 0.6% |
| Base accuray - Quality Value (kmer) |  | 32.8 | 44.5 |
| Number of genes |  | 109,552 | 107,891 |
| Average number of exons |  | 5.10 | 5.33 |
| BUSCO on gene predictions<br>(N=4,896) | Complete | 99.1% | 99.5% |
|  | Duplicated | 94.6% | 98.2% |
|  | Fragmented | 0.2% | 0.1% |
|  | Missing | 0.7% | 0.4% |

The completeness and quality of the assembly was estimated by searching for the presence of known genes, i.e. the 107,891 High Confidence (HC) genes predicted in CS RefSeq v1.1. We used BLAST[25] to search for the presence of each of the 461,476 exons larger than 30 bps in the Renan scaffolds, and we considered only matches showing at least 95% identity over at least 95% query length. We found hits for 96.2% of the query exons with on average 99.3% identity, suggesting that the gene space is assembled at a high-quality level. The missing genes/exons would correspond, in most of the cases, to real presence/absence variations between CS and Renan while the nucleotide divergence between exons is 0.7%. It was the first evidence that homoeologous gene copies, sharing on average 97% identity[7], were not collapsed in the Renan assembly. We confirmed this by showing that 62% of the CS exons are strictly identical in Renan (and carried by the same chromosome). Such level of nucleotide divergence between CS and Renan is similar to what has been shown through whole genome alignments (Brinton et al. 2020).

We then assessed the assembly quality of the TE space by aligning the complete dataset of ISBP markers of CS onto the Renan assembly. We found that 94% markers were conserved (at least 90% identity over 90% query length) i.e., present in the assembly, revealing that the TE space is extremely close to completeness. Indeed, 6% of missing markers is similar to the proportion of expected Presence-Absence variations (PAVs) affecting TEs[26].

Additionally, we searched for telomeric repeats (TTTAGGG) in the 21 chromosomes and found telomeric repeats at both ends of chromosome 7A, which is generally an indicator of the completion of the chromosome sequence. Both ends of chromosome 7A were also validated by the optical map (Figure S1).

### Impact of the polishing

Based on BUSCO and the alignment of the IBSP markers from the CS assembly, we monitored the evolution of the consensus quality through successive polishing iterations. As previously described, the SMARTdenovo consensus allowed the recovery of a greater number of complete BUSCO genes compared to that of Flye, which may be an indicator of its greater accuracy. However, the BUSCO score was still low (83%) especially for a hexaploid genome, underlining the importance of polishing raw assemblies. Likewise, we were able to find 80.4% of the IBSP markers but only 7% were aligned without mismatch between the two genotypes (Table S5). When polished with long-reads, the BUSCO score reached 96.7% and 92.9% of the IBSP markers were retrieved (including 28.0% with perfect matches). The subsequent polishing step with short reads weakly decreased the BUSCO score (from 96.7% to 96.6%), but the proportion of duplicated genes increased from 83.1% to 87.0% which is here wanted because in the case of a hexaploid genome most of the genes are in three copies. Moreover, the proportion of perfectly aligned ISBP markers drastically increased from 28.0% up to 58.9%. Although the polishing with short reads weakly impacts the BUSCO conserved genes, the IBSP markers underline its importance in the case of long reads assemblies. Since ISBPs are unique tags sampling the whole genome, this analysis revealed that nucleotide errors were frequent before polishing, affecting half of the sample loci. Thus, we showed that the polishing steps were successful, even in this large and polyploid genome, and drastically improved the quality of the consensus.

### Recent improvement of the ONT technology

Oxford Nanopore Technology is evolving rapidly, and improvements to the base calling softwares are frequent, allowing old data to be analyzed with the aim of improving read accuracy and subsequent analysis. To measure the gain brought by each new version during this project, we analyzed a subset of ultra-long reads (longer than 100kb) with different basecallers or versions of the same basecaller: guppy 2.0, guppy 3.0.3 (High Accuracy mode), guppy 3.6 (High Accuracy mode) and the recent bonito v0.3.1. We observed a strong difference in accuracy, of around 7%, between guppy 2.0 and the newer basecaller (bonito v0.3.1), representing the gain over the last two years (Figure S2A). This significant improvement could lead nanopore users to reanalyze their old sequencing data to improve the quality of their assemblies. As an example, the accuracy of raw nanopore reads gained about 2% on average using guppy 3.6 (Table S6). We observed a reduction of the number of contigs of 19%, and an improvement of the contig N50 of 26%. Likewise, the cumulative size is slightly higher in the guppy 3.6 assembly, which may underline a smaller amount of collapsed repetitive regions (Table S7).

More importantly, the identity percentage obtained when aligning ONT reads on the wheat assembly is lower than what was obtained on yeast and human samples (Figure S2B). This difference can be explained by the fact that, first, the consensus of the wheat genome is not perfect and secondly, that basecallers are trained on a mixture which contains yeast and human data. Indeed, DNA modification patterns can differ between taxa, and read accuracy seems better when the model was trained on native DNA from the same species[27]. This huge difference between the read accuracy of yeast and wheat samples should motivate nanopore users to train basecaller models to their targeted species.

### Annotation of transposable elements and protein-coding genes

We annotated TEs based on similarity search against our wheat-specific TE library ClariTeRep[28] and raw results were then refined using CLARITE, a homemade program able to resolve prediction conflicts, merge adjacent features into a single complete element, and identify nested insertion patterns. We detected 3.9 million copies of TEs in the Renan genome assembly, representing 12.0 Gb i.e. 84% of the assembly size. The proportions of each superfamily were similar to what has been described for CS[29] (Table 2). Gene annotation was achieved by, first, transferring genes predicted in CS RefSeq v2.1 by homology using the MAGATT pipeline[22]. This allowed us to accurately transfer 105,243 (out of 106,801; 98%) HC genes and 155,021 (out of 159,846; 97%) Low Confidence genes. Such a transfer of genes predicted in another genotype (here CS) avoided genome-wide *de novo* gene prediction that may artificially lead to many differences between the annotations. We thus focused *de novo* predictions using TriAnnot[30] only on the unannotated part of the genome, representing 8.5% of the 14.2 Gb, after having masked transferred genes and predicted TEs. For that purpose, we produced RNASeq data for Renan from 28 samples corresponding to 14 different organs/conditions in replicates: grains at four developmental stages (100, 250, 500, and 700 degree days) under heat stress and control conditions, stems at two developmental stages, leaves at three stages, and roots at one stage), representing on average 78.8 million read-pairs per sample i.e 2.2 billion read-pairs in total. This method allowed us to predict 4,440 genes specific to Renan compared to CS i.e., 4% of the gene complement. This is consistent with the extent of structural variations affecting genomes of *Triticeae*[26]. Transfer of known genes, novel predictions, and manual curation (limited to storage protein encoding genes), led us to annotate 109,552 protein-coding genes on the Renan pseudomolecules.

**Table 2:** TE classes proportions in Chinese Spring and Renan genome assemblies.

|  | Chinese Spring<br>RefSeq_v1.0<br>from Zhu et al. [22] | Chinese Spring<br>RefSeq_v2.1<br>from Zhu et al.[22] | Renan RefSeq_v2.0 |
| --- | --- | --- | --- |
| Genome size (bp) | 14,066,280,851 | 14,225,829,371 | 14,195,643,615 |
| TE (bp) | 11,921,309,743 | 12,092,094,168 | 11,967,447,100 |
| TE (%) | 84.7 | 85.0 | 84.3 |
| **Class I (Retrotransposons)** | **67.6** | **66.9** | **66.6** |
| Gypsy (RLG) | 46.7 | 46.1 | 45.8 |
| Copia (RLC) | 16.7 | 16.5 | 16.5 |
| Unclassified LTR<br>retrotransposons (RLX) | 3.24 | 3.3 | 3.2 |
| LINE (RIX) | 0.9 | 1.1 | 1.1 |
| SINE (SIX) | 0.01 | 0.01 | 0.01 |
| **Class II (DNA transposons)-**<br>**Subclass 1** | **16.5** | **17.0** | **16.9** |
| CACTA (DTC) | 15.5 | 15.9 | 15.8 |
| Mutator (DTM) | 0.38 | 0.44 | 0.44 |
| Unclassified DNA<br>transposons with TIR (DTX) | 0.21 | 0.24 | 0.24 |
| Harbinger (DTH) | 0.16 | 0.18 | 0.18 |
| Mariner (DTT) | 0.16 | 0.17 | 0.17 |
| Unclassified DNA<br>transposons (DXX) | 0.06 | 0.06 | 0.06 |
| hAT (DTA) | 0.006 | 0.009 | 0.009 |
| Helitrons (DHH) | 0.004 | 0.01 | 0.01 |
| **Unclassified TE (XXX)** | **0.68** | **0.95** | **0.82** |

### Comparison with existing hexaploid genome assemblies

We compared our long-read assembly with 10 other available chromosome-scale assemblies of wheat genomes. Although the gene content was similar between the different assemblies, as expected, the assemblies based on short reads had a lower contiguity (contig N50 values lower than 100kb compared to the 2 Mb of the assembly of the Renan genome, Figure 2A-B). Logically, they also contained more gaps (around 40 times, Figure 2C). Interestingly, we found more gaps per Mb in the D subgenome compared to the A and B subgenomes in Renan (Figure S3). This indicates that the D subgenome is more difficult to assemble even though it has a smaller genome size and contains less repetitive elements. The same trend was already observed in another polyploid genome, the rapeseed and its two subgenomes A and C[11]. Chromosomes from the different assemblies had similar length except for the *ArinaLrFor* and the SY_Mattis variety in which a translocation has been previously described between chromosomes 5B and 7B[8] (Figure 2D).

**Figure 2.**
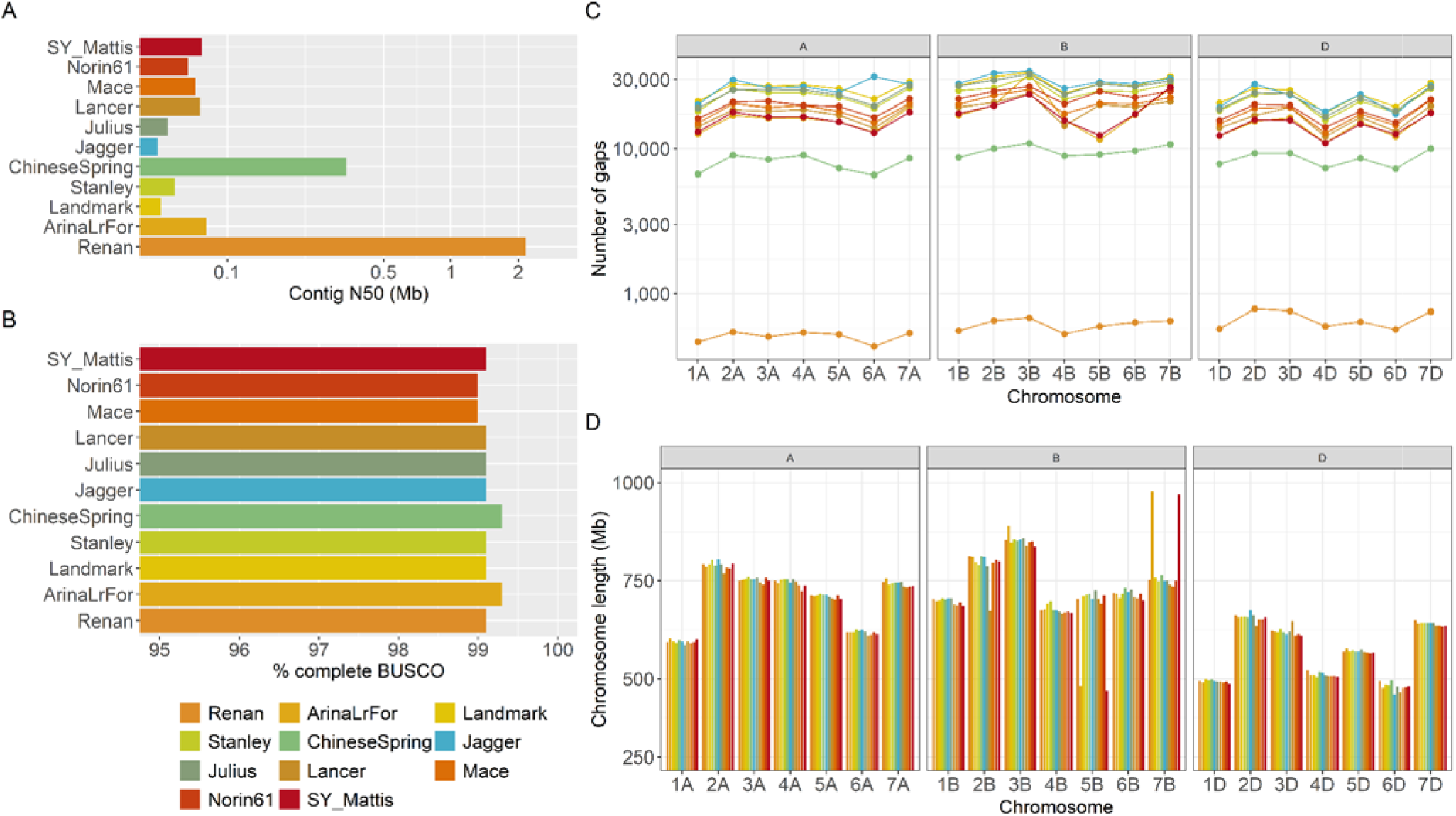
Comparison of existing hexaploid genome assemblies **A.** contig N50 values in Mbp. **B.** Proportion of complete BUSCO genes found in each assembly (N=4,896). **C.** Number of gaps in each chromosome. **D.** chromosome length in Mb.

In addition, we generated dotplots between CS and Renan homeologous chromosomes and confirmed the strong collinearity between the two genomes (Figure 3). Whole chromosome alignments highlighted 16 large-scale inversions (>5 Mb; up to 118 Mb) on 10 chromosomes and 1 translocation of a ca. 45 Mb segment on chromosome 4A. We performed the same comparisons with the 10 other available genomes of related varieties assembled at the pseudomolecule level (Supplementary Data 1). It showed that only 2 of these inversions are specific to Renan while the others are shared between several accessions. They correspond to regions of 23 Mb on chr6B (position 398-421 Mb) and 10 Mb on chr7B (position 267-277 Mb).

**Figure 3.**
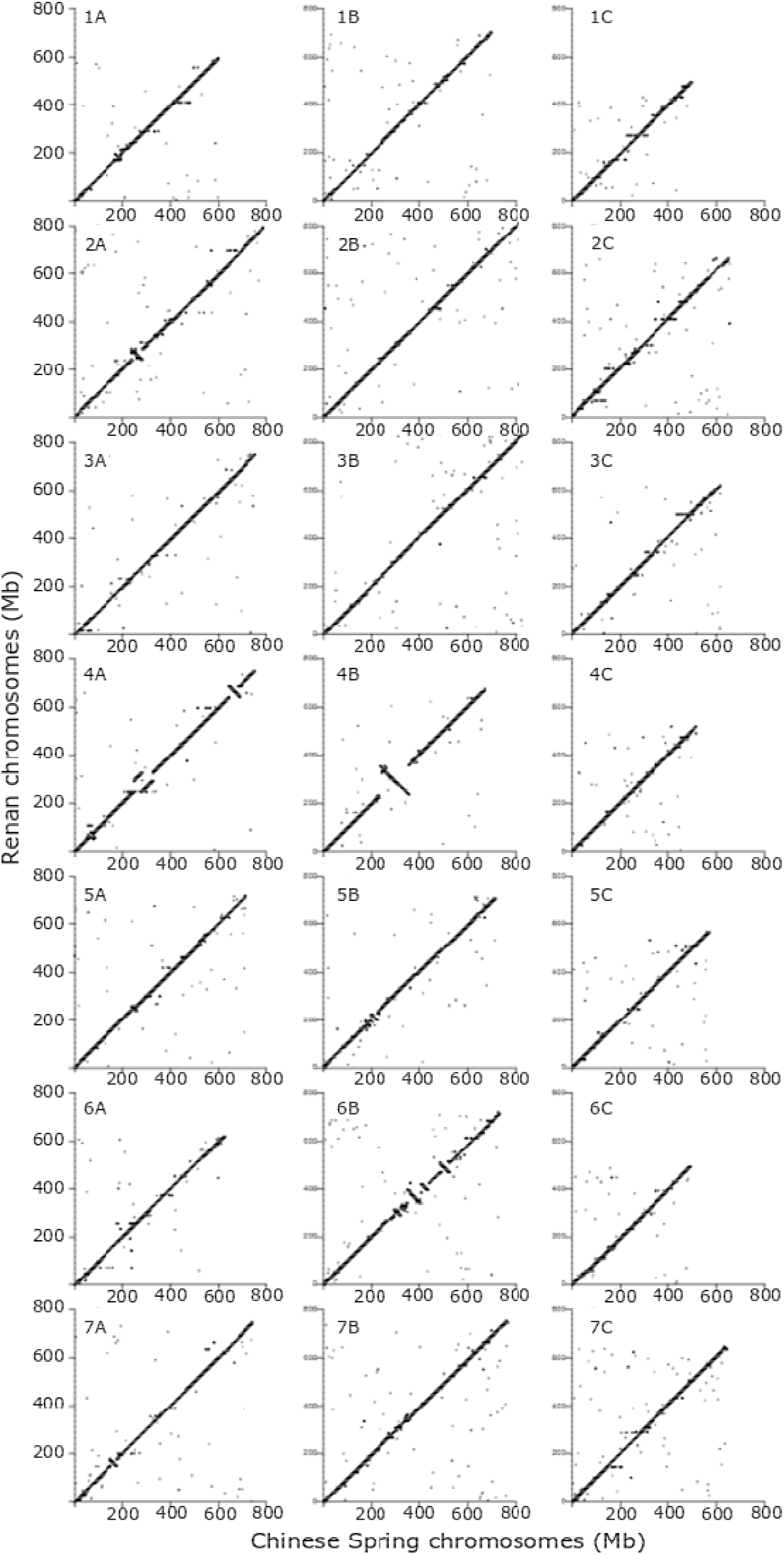
Dotplot comparisons of the 21 chromosomes of Renan (y axis) with the Chinese Spring RefSeq v2.1 assembly (x axis).

### Haplotype characterisation

Crop breeding involves the selection of desired traits and their combination to generate improved genotypes. Generally, these traits correspond to genomic regions carrying genetic variations or genes[31]. These regions of interest are inherited from their parents in the form of large genomic blocks. The availability of several assemblies of the wheat genome now allows the detection of these haplotypic blocks. Using the 11 chromosome-scale wheat assemblies and an approach based on colored de Bruijn graphs, we investigated these haplotypic blocks and applied our method to the 21 chromosomes of wheat. First, a colored de Bruijn graph was built for each chromosome, where each colour represents a different cultivar. Short (1kb) and evenly distributed (every 20kb) markers were extracted from each chromosome and compared to the colored de Bruijn graph to extract their presence/absence in each wheat cultivar. On each chromosome, the 15 most abundant presence/absence profiles were selected and used to characterise haplotypic blocks. The haplotype blocks of chromosome 6A, which is associated with productivity traits (as for example yield, grain size and height), have already been expertized using a different method[31]. We obtained similar results (Figure 4), except for the Chinese Spring chromosome 6A. Previous results have assigned a unique haplotype to this wheat line. But in our case Chinese Spring exhibits the same haplotype as SY Mattis, Jagger, Lancer and Norin61, which had previously been described as sharing the same haplotype. These differences may be explained by the stringency of the comparison, which perhaps should be adjusted separately for each chromosome. Concerning the Renan cultivar, the chromosome 6A has haplotype blocks similar to those of the ArinaLrFor line. Additionally, we used this method to investigate haplotypic blocks that are specific to one or a subset of wheat cultivars.

**Figure 4.**
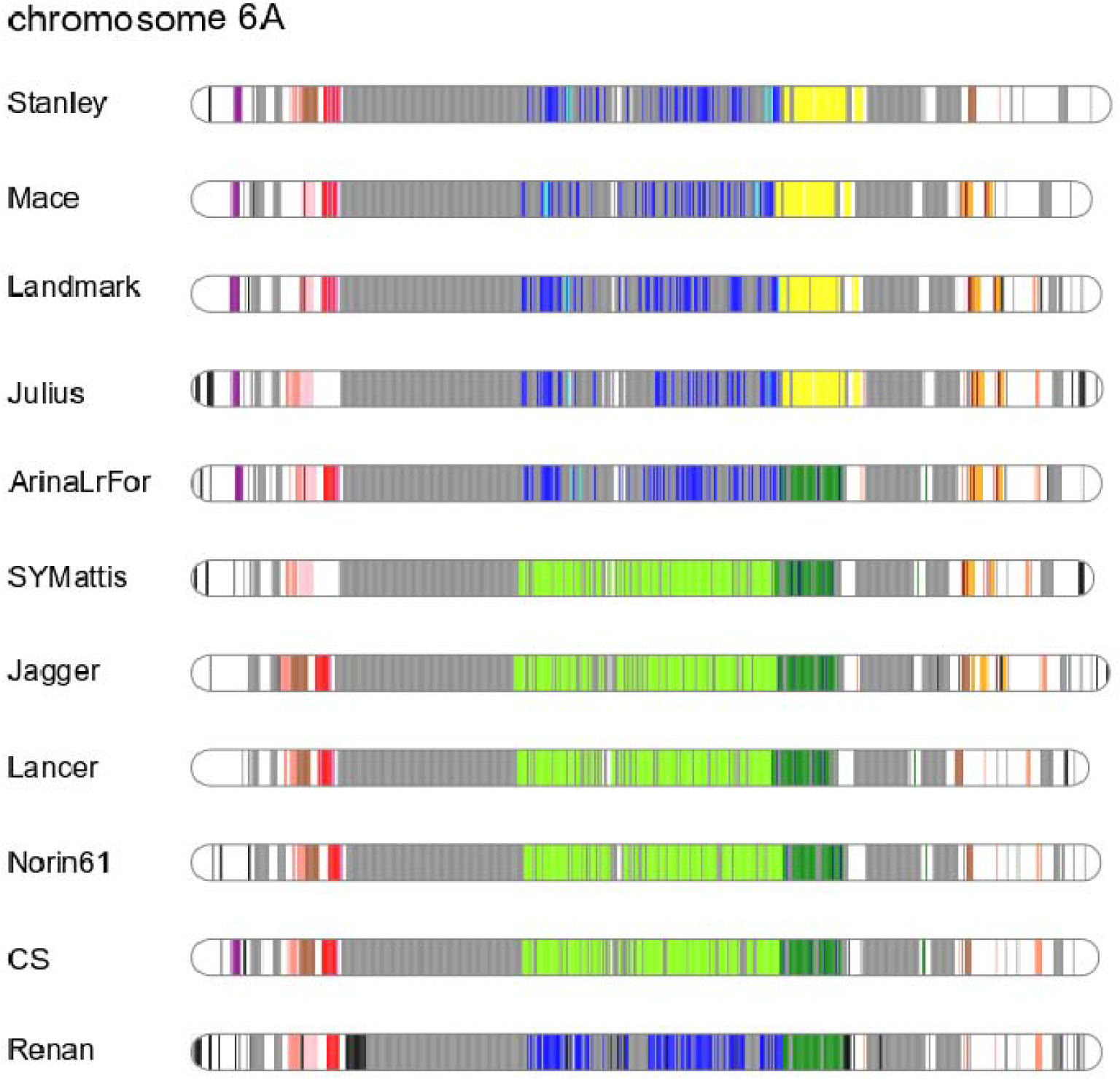
Representation of haplotype blocks in chromosome 6A for the 11 chromosome-scale cultivars (based on 1-Mbp blocks). Regions with the same colour represent common regions in wheat lines, except white regions which are not contained in haplotype blocks. The gray and black regions represent haplotypes respectively shared by at least 10 cultivars or specific to a given cultivar.

### Identification of introgressions

Introgression is an important source of genetic variation which is generally the signature of breeding programmes, especially in wheat[32]. Several introgressions have already been reported[8], notably in chromosomes 2B and 3D in LongReach Lancer and in chromosome 2A in Jagger, Mace, SY Mattis and CDC Stanley. Using our approach, we were able to clearly identify the two introgressions in LongRead Lancer (Figure 5a), and the *Ae. ventricosa* introgression in chromosome 2A (Figure 5b). In addition, we found that this introgression of *Ae. ventricosa* is also present in the Renan cultivar (Figure 5b). The optical map was aligned with this 34 Mb region of Renan and validated the correct structure of this important region carrying multiple resistance genes (Yr17, Lr37, Sr38, Cre5). More importantly, the 34 Mb consisted of 22 contigs in Renan and 2,339 in Jagger. A comparison of the fragmentation near the introgression point is presented in Figure 5d and shows a large difference between the long- and short-reads assemblies. Additionally, we also identified several candidate introgressions, which had already been spotted through retrotransposon profiles[8]: i) a 45-Mb region on chromosome 2D which is shared between the lines Julius, ArinaLrFor, SY Mattis, Jagger and also Renan (Figure 6a); ii) a 53-Mb region at the end of chromosome 3D in Lancer (Figure 6b); iii) a 48-Mb region at the beginning of chromosome 3D in SY Mattis (Figure 6b) and iv) the *Ae. ventricosa* introgression of 30-Mb in chromosome 7D which carries Pch1 resistance gene (Figures 6c).

**Figure 5.**
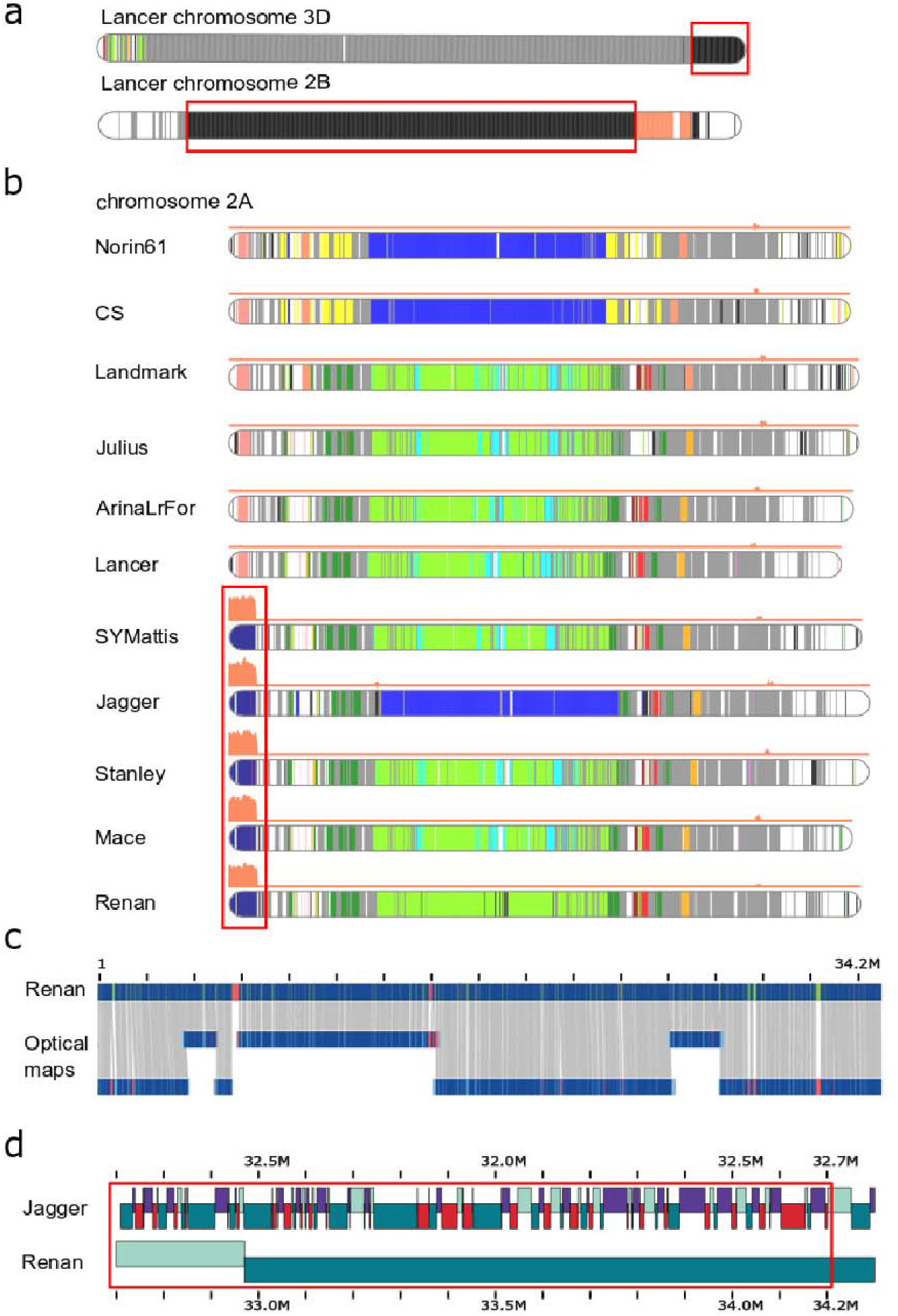
Haplotypic blocks in wheat chromosomes. Colors represent common regions in wheat cultivars. The gray and black regions represent haplotypes respectively shared by at least 10 cultivars or specific to a given cultivar. The orange curve, when present, represents coverage with *Ae. ventricosa* short reads. The red boxes frame the introgressions. **a.** Known introgressions in chromosomes 3D and 2B in Lancer. Regions in black represent genomic regions that are specific to Lancer and are respectively *T. ponticum* and *T. timopheevii* introgressions as described previously[8]. **b.** *Ae. ventricosa* introgression on chromosome 3D in Stanley, Mace, SY Mattis and Jagger. This known introgression is also present in Renan. The dark blue block represents the region shared across the five cultivars. **c.** Validation of the introgression in Renan (chromosome 2A from 1 to 34.2Mb) using Bionano maps. **d.** Comparison of the contig composition of the first megabases from the introgression point in Jagger and Renan cultivars.

**Figure 6.**
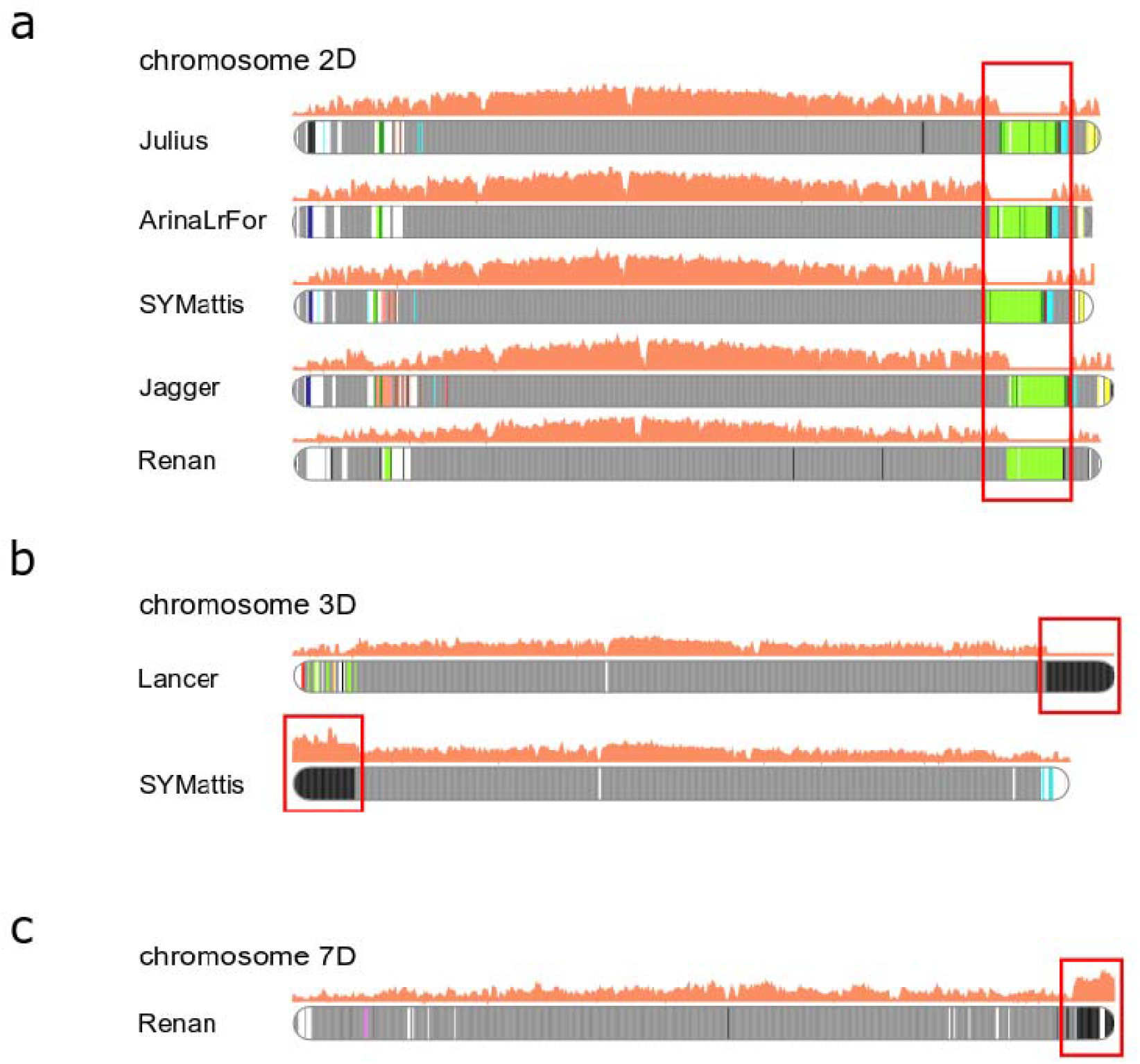
Haplotypic blocks in wheat chromosomes. Colors represent common regions in wheat cultivars. The gray and black regions represent haplotypes respectively shared by at least 10 cultivars or specific to a given cultivar. The orange curve represents coverage with *Ae. ventricosa* short reads. The red boxes frame the introgressions. **a.** Candidate introgression (green block) on chromosomes 2D in Julius, ArinaLrFor, SY Mattis, Jagger and Renan. **b.** Candidate introgressions (black blocks) on chromosome 3D in Lancer and SY Mattis. **c.** *Ae. ventricosa* introgression (black block) on chromosome 7D in Renan.

**Figure 7.**
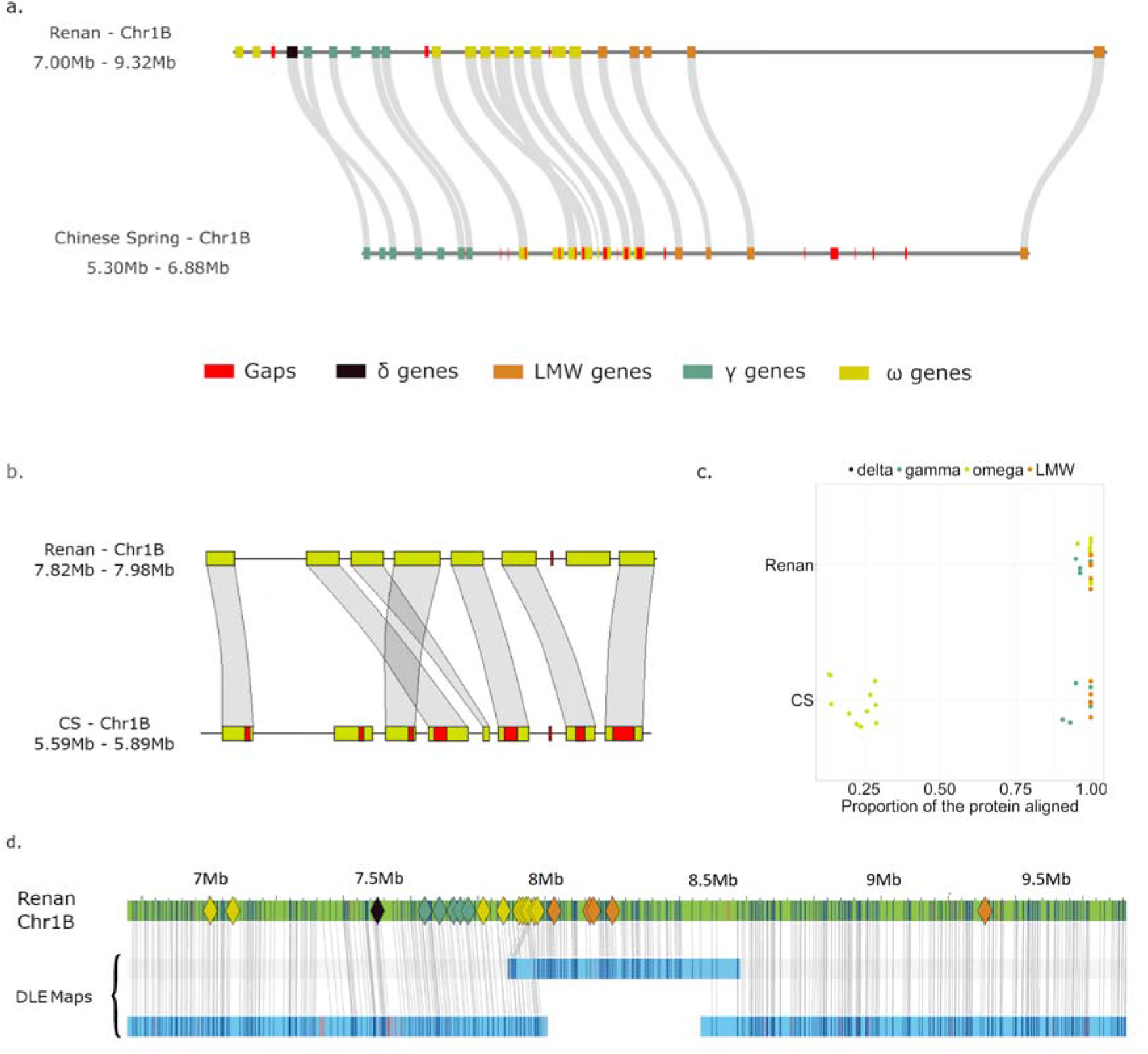
Comparative view of an important locus on chromosome 1B containing prolamin and resistance genes, tandemly duplicated. **a.** Representation of the region with gaps and genes on the two assemblies of Renan and CS. **b.** Zoomed view on the omega gliadin gene cluster **c.** Proportion of the length of the proteins that were aligned in the genomic region of Renan and CS. **d.** Alignment view of Bionano maps on the Renan cluster, colored diamond shapes represent genes belonging to the omega gliadin gene cluster. The optical maps are in blue and the chromosome sequence in green. Restriction sites are represented by vertical lines and are joined between the sequence and the map when properly aligned.

Moreover, a known large-scale structural variation in chromosomes 5B and 7B of ArinaLrFor and SY Mattis cultivars was also easily identifiable using haplotypic blocks of individual chromosomes (Figure S4).

### Comparative analysis of a storage protein coding gene cluster in *T. aestivum*

Tandem duplications are an important mechanism in plant genome evolution and adaptation[33,34] but the assembly of tandemly duplicated gene clusters is difficult, especially with short reads. In order to illustrate the gain brought by this optimized assembly process, we focused on an important locus on chromosome 1B known to carry multiple copies of storage protein and disease resistance genes[35,36]. Among them, the genes encoding omega-gliadins are not only duplicated in tandem, but are also composed of microsatellite DNA in their coding part, making them particularly hard to assemble properly from short reads. We compared orthologous regions harboring these genes between CS and Renan, spanning 1.58 Mb and 2.32 Mb, respectively. The CS region was more fragmented with 101 gaps versus only 3 in Renan (Figures 5a). The number of copies of omega-gliadin encoding genes was quite similar: 9 in CS and 10 in Renan. The most striking difference came from the completeness of the microsatellite motifs: 8 copies out of 9 contain N stretches in CS RefSeq v2.1, revealing that the microsatellite is usually too large to be fully assembled with short reads (Figure 5b). In contrast, all 10 copies predicted in Renan were assembled completely. More generally, we mapped the corresponding proteins back to the locus and showed that it was better reconstructed in the Renan assembly, with a mean protein alignment length of 99% compared to 58% in CS (Figure 5c). In addition, the optical map was used to validate the structure of this region in Renan and the assembly was consistent with the three maps of this loci (Figure 5d).

### Comparative analysis of the locus that provides resistance to the orange wheat blossom midge

Like a few other wheat cultivars, Renan is resistant to the orange wheat blossom midge (OWBM). The *Sm1* gene is known to confer resistance to wheat and a previous study has shown that CDC Landmark is also resistant to the OWBM, and carries a 7.3-Mb haplotype within the *Sm1* locus on chromosome 2B[8]. We extracted and aligned the corresponding region of CDC Landmark on each cultivar, to precisely locate the corresponding region on each chromosome 2B. From these eleven regions of 1-2 Mb, we computed the haplotypic blocks using a higher resolution than previously (1 kb marker every 5 kb). This analysis revealed a strong similarity of the *Sm1* locus between CDC Landmark and Renan (Figure 8a), the presence of the *Sm1* gene in blocks shared between the two cultivars. In addition, a comparison of the fragmentation of these two regions underlines the higher contiguity of the Renan assembly, with 4 contigs in the Renan *Sm1* locus compared to 62 in CDC Landmark (Figure 8b). The *Sm1* locus of Renan is in agreement with the optical map and shows clearly the three remaining gaps that may correspond to smaller and unanchored contigs.

**Figure 8.**
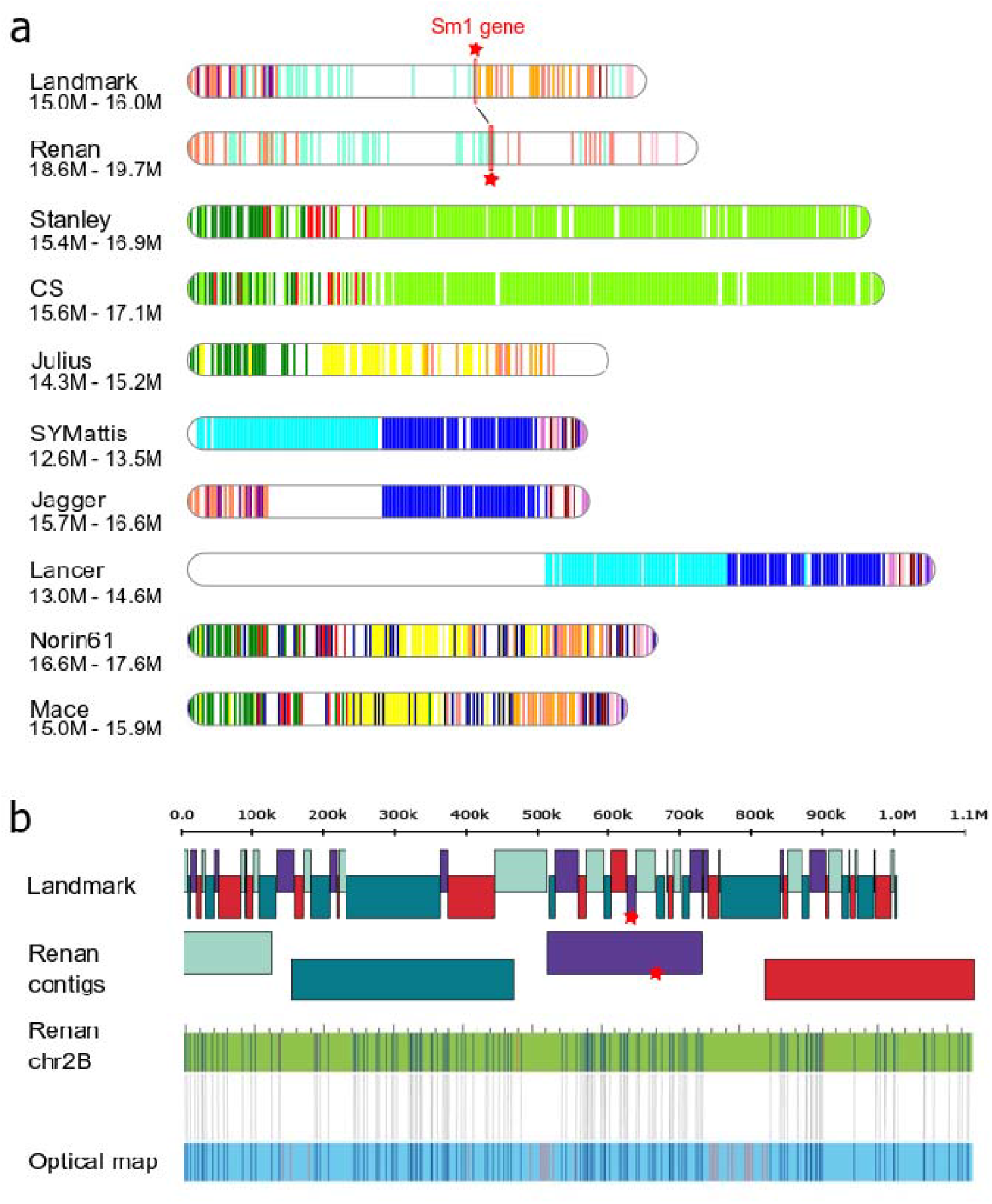
Comparison of the *Sm1* loci. **a.** Representation of haplotype blocks (5kb bins) of the region surrounding the *Sm1* gene on chromosome 2B. Colors represent common regions in wheat cultivars. The genomic region of Landmark (15Mb to 16Mb) was aligned against other cultivars to localize the *Sm1* loci. The *Sm1* gene in Landmark and Renan, the two *Sm1* carrier cultivars, is represented by a red star. **b.** Comparison of the contig composition in the *Sm1* region of Landmark and Renan, and validation of the assembly structure in Renan using Bionano optical maps. The optical map is in blue and the chromosome sequence in green. Restriction sites are represented by vertical lines and are joined between the sequence and the map when properly aligned..

## Discussion

In this study, we showed that the recent improvement of the Oxford Nanopore Technology, in terms of error rate and throughput, has opened up new perspectives in the age of long-read technologies. Indeed, the sequencing and assembly of complex genomes, like the hexaploid wheat, is now accessible to sequencing facilities. Additionally, the ability to sequence ultra-long reads using ONT devices is a real advantage over the other long-read technology, namely PACBIO. In this study, we were able to generate a coverage of 14X with reads longer than 50kb, whereas PACBIO libraries, used to generate HiFi (High-Fidelity) reads, are generally sized around 15kb[37,38]. Several studies have already underlined the positive impact of these ONT ultra-long reads on the assembly contiguity[9,37,39]. In contrast, the error rate that was previously a thorn in their side has been drastically reduced over the last year. Herein we reported a quality score near Q10-Q15 for individual ONT reads, as already shown[27], which is still far from what HiFi reads can provide, generally near Q30[37]. The high accuracy of HiFi reads might be sufficient to distinguish copies from repeat regions if they present few variations. The impact of ultra-long reads will lie mainly in the case of identical repeats, and obviously, the presence of these particular cases will depend on the evolutionary history of the studied genomes. In addition, this high error rate has an impact on the consensus quality, and at the moment, a combination of ONT and Illumina reads is still needed to achieve a decent per-base accuracy.

By following basecallers evolution, we noticed that the gain when using recent basecaller is high and we guess this observation will encourage users to reprocess older data. However, this is not trivial and it requires sufficient computing resources. Interestingly, we observed that the error rate of ONT data is organism dependent and that the training of basecaller has a significant impact on the overall quality of the reads[27]. This is, in our opinion, an important fact because a large proportion of *de novo* assemblies now concern non-model organisms and users will have to address this limitation of current software. There are existing methods to train the basecaller on non-model species[40,41], but this can still be a big barrier, depending on the size of the dataset, for many end users. However, as highlighted in this study, the combination of long- and short-reads sequencing with polishing methods greatly improves the consensus sequence of a given genome assembly and these algorithms seem sufficient at least in coding regions.

Even though there are now several chromosome-scale assemblies of the hexaploid wheat genome, this assembly of the Renan variety based on long-reads will benefit biologists and geneticists as it offers a high resolution. We show that our chromosome-scale assembly of Renan based on long reads can bring new insight into genomic regions of interest. In particular, in regions that carry multiple resistance genes, as a large *Ae. ventricosa* introgression shared with other cultivars on chromosome 2A and a unique *Ae. ventricosa* introgression on chromosome 7D. The lower number of gaps in these regions will help to localize genes of interest and to have a better understanding of the impact of these introgressions. Additionally, we demonstrated by examining two important locus, containing prolamin and resistance genes that such regions are truly enhanced and contain very few gaps compared to assemblies based on short reads.

Moreover, unlike recent chromosome-scale assemblies, Renan’s gene prediction is not only a projection of Chinese Spring gene models, but also includes *de novo* annotation with RNA-Seq data which is of real benefit for the construction of pan genome (or pan annotation) or when cultivar-specific genes are examined. For all of these reasons, we believe this high resolution assembly will benefit the wheat community and help breeding programs dedicated to the bread wheat genome.

## Methods

### Plant material and DNA extraction

*Triticum aestivum* cv. Renan seeds were provided by the INRAE Biological Resource Center on small grain cereals and grown for two weeks and a dark treatment was applied on the seedlings for two days before collecting leaf tissues.

For the sequencing experiments, DNA was isolated from frozen leaves using QIAGEN Genomic-tips 100/G kit (Cat No./ID: 10243) and following the tissue protocol extraction. Briefly, 1g of leaves were ground in liquid nitrogen with mortar and pestle. After 3h of lysis and one centrifugation step, the DNA was immobilized on the column. After several washing steps, DNA is eluted from the column, then desalted and concentrated by alcohol precipitation. The DNA is resuspended in the TE buffer.

To generate the optical map, uHMW DNA were purified from 0.5 gram of very young fresh leaves according to the Bionano Prep Plant tissue DNA Isolation Base Protocol (30068 - Bionano Genomics) with the following specifications and modifications. Briefly, the leaves were fixed using a fixing solution (Bionano Genomics) containing formaldehyde (Sigma-Aldrich) and then grinded in a homogenization buffer (Bionano Genomics) using a Tissue Ruptor grinder (Qiagen). Nuclei were washed and embedded in agarose plugs. After overnight proteinase K digestion in Lysis Buffer (Bionano Genomics) and one hour treatment with RNAse A (Qiagen), plugs were washed four times in 1x Wash Buffer (Bionano Genomics) and five times in 1x TE Buffer (ThermoFisher Scientific). Then, plugs were melted two minutes at 70°C and solubilized with 2 μL of 0.5 U/μL AGARase enzyme (ThermoFisher Scientific) for 45 minutes at 43°C. A dialysis step was performed in 1x TE Buffer (ThermoFisher Scientific) for 45 minutes to purify DNA from any residues. The DNA samples were quantified by using the Qubit dsDNA BR Assay (Invitrogen). Quality of megabase size DNA was validated by pulsed field gel electrophoresis (PFGE).

### Illumina Sequencing

DNA (1.5μg) was sonicated using a Covaris E220 sonicator (Covaris, Woburn, MA, USA). Fragments (1μg) were end-repaired, 3’-adenylated and Illumina adapters (Bioo Scientific, Austin, TX, USA) were then added using the Kapa Hyper Prep Kit (KapaBiosystems, Wilmington, MA, USA). Ligation products were purified with AMPure XP beads (Beckman Coulter Genomics, Danvers, MA, USA). Libraries were then quantified by qPCR using the KAPA Library Quantification Kit for Illumina Libraries (KapaBiosystems), and library profiles were assessed using a DNA High Sensitivity LabChip kit on an Agilent Bioanalyzer (Agilent Technologies, Santa Clara, CA, USA). The library was sequenced on an Illumina NovaSeq instrument (Illumina, San Diego, CA, USA) using 150 base-length read chemistry in a paired-end mode. After the Illumina sequencing, an in-house quality control process was applied to the reads that passed the Illumina quality filters[42]. These trimming and removal steps were achieved using Fastxtend tools (https://www.genoscope.cns.fr/fastxtend/).

### Nanopore Sequencing

Libraries were prepared according to the protocol Genomic DNA by ligation (SQK-LSK109 kit). Genomic DNA fragments (1.5 μg) were repaired and 3’-adenylated with the NEBNext FFPE DNA Repair Mix and the NEBNext^®^ Ultra™ II End Repair/dA-Tailing Module (New England Biolabs, Ipswich, MA, USA). Sequencing adapters provided by Oxford Nanopore Technologies (Oxford Nanopore Technologies Ltd, Oxford, UK) were then ligated using the NEBNext Quick Ligation Module (NEB). After purification with AMPure XP beads (Beckmann Coulter, Brea, CA, USA), the library was mixed with the Sequencing Buffer (ONT) and the Loading Bead (ONT) and loaded on MinION or PromethION R9.4.1 flow cells. One PromethION run was performed with Genomic DNA purified with Short Read Eliminator kit (Circulomics, Baltimore, MD, USA) before the library preparation.

### Optical Maps

Labeling and staining of the uHMW DNA were performed according to the Bionano Prep Direct Label and Stain (DLS) protocol (30206 - Bionano Genomics). Briefly, labeling was performed by incubating 750 ng genomic DNA with 1× DLE-1 Enzyme (Bionano Genomics) for 2 hours in the presence of 1× DL-Green (Bionano Genomics) and 1× DLE-1 Buffer (Bionano Genomics). Following proteinase K digestion and DL-Green cleanup, the DNA backbone was stained by mixing the labeled DNA with DNA Stain solution (Bionano Genomics) in presence of 1× Flow Buffer (Bionano Genomics) and 1× DTT (Bionano Genomics), and incubating overnight at room temperature. The DLS DNA concentration was measured with the Qubit dsDNA HS Assay (Invitrogen).

Labelled and stained DNA was loaded on Saphyr chips. Loading of the chips and running of the Bionano Genomics Saphyr System were all performed according to the Saphyr System User Guide (30247 - Bionano Genomics). Data processing was performed using the Bionano Genomics Access software.

A total of 4541 Gb data were generated. From this data, molecules with a size larger than 150kb were filtered generating 1931 Gb of data. These filtered data, corresponding to 128x coverage of the *Triticum aestivum* cv. Renan consists of 7,810,298 molecules with an N50 of 237.5kb and an average label density of 14.3/100kb. The filtered molecules were aligned using RefAligner with default parameters. It produced 1053 genome maps with a N50 of 37.5 Mbp for a total genome map length of 14946.8 Mbp.

### RNA extraction

Several tissues (stem, leaves, root or grain) were collected on plants with different growth conditions and of different ages. Each of these 28 tissues was subjected to RNA extraction with the following protocole: 200mg to 1g of fine powder was put in a 50ml falcon tube with 4.5 ml of NTES buffer [0.1 M NaCl, 1% SDS, 10 mM Tris-HCl (pH 7.4), 1 mM EDTA(pH 8)]. After vortexing the tube, 3ml of phenol-chloroforme-IAA were added. The tube was mixed for 10 minutes and centrifuged for 20 minutes at 5,000 rpm (15°C). The aqueous phase was collected and placed in a new 15ml tube. 3ml of phenol-chloroforme-IAA were added. The tube was mixed for 10 minutes and centrifuged for 20 minutes at 5,000 rpm (15°C). The aqueous phase was collected and placed in a new 50ml tube. 1/10 of AcNa 3M (pH 5.2) and 2 volumes of 100% ethanol were added. The tube was mixed gently by turning and centrifuged 20 minutes at 5,000 rpm (4°C). The supernatant was removed. The precipitate was dried and resuspended in 20□μl RNAse free water. A treatment with DNase was realized and the RNA were purified on a MinElute column (Qiagen). A second treatment with DNAse was realized by adding DNAse directly on the filter. After ethanol cleanup, the column was eluted with 14 μl of RNAse free water. The quality of the RNA was evaluated using RNA 6000 Nano Assay chip for size and RIN estimation and spectrophotometry (A260/A280 and A260/A230 ratios) for purity estimation. The RNA were quantified using Qubit RNA high sensitivity Assay kit (Invitrogen).

### RNA sequencing

RNA-Seq library preparations were carried out from 500ng to 2000ng of total RNA using the TruSeq Stranded mRNA kit (Illumina, San Diego, CA, USA), which allows mRNA strand orientation (sequence reads occur in the same orientation as antisense RNA). Briefly, poly(A)+ RNA was selected with oligo(dT) beads, chemically fragmented and converted into single-stranded cDNA using random hexamer priming. Then, the second strand was generated to create double-stranded cDNA. cDNA were then 3’-adenylated, and Illumina adapters were added. Ligation products were PCR-amplified. Ready-to-sequence Illumina libraries were then quantified by qPCR using the KAPA Library Quantification Kit for Illumina Libraries (KapaBiosystems, Wilmington, MA, USA), and libraries profiles evaluated with an Agilent 2100 Bioanalyzer (Agilent Technologies, Santa Clara, CA, USA). Each library was sequenced using 151 bp paired end reads chemistry on an Illumina NovaSeq 6000 sequencer (Illumina, San Diego, CA, USA).

### Long reads genome assembly

The 20 ONT runs were basecalled using two versions of guppy: 3.3 HAC and 3.6 HAC (Table S6). We monitored the gain of each guppy basecaller release and evaluated three different assemblers in the context of large genomes: Redbean[16] v2.5 (git commit 3d51d7e), SMARTdenovo[17] (git commit 5cc1356) and Flye[18] v2.7 (git commit 5c12b69). All assemblers were launched using a subset of reads consisting of 30X of the longest reads (Table S3). Then, we selected one of the assemblies based not only on contiguity metrics such as N50 but also cumulative size, proportion of unknown bases. The Flye (longest reads) and SMARTdenovo (all reads) assemblies were very similar in terms of contiguity but we decided to keep the SMARTdenovo assembly as its cumulative size was higher. The SMARTdenovo assembler using the longest reads resulted in a contig N50 of 1.1Mb and a cumulative size of 14.07Gb. As nanopore reads contain systematic error in homopolymeric regions, we polished the consensus of the selected assembly with nanopore reads as input to the Racon (v1.3.2, git commit 5e2ecb7) and Medaka softwares. In addition, we polished the assembly two additional times using Illumina reads as input to the Hapo-G tool (v1.0, git commit).

### Long range genome assembly

The Bionano Genomics scaffolding workflow (Bionano Solve version 3.5.1) was launched with the nanopore contigs and the Bionano map. We found in several cases that the nanopore contigs were overlapping (based on the optical map) and these overlaps were corrected using the BiSCoT software[21] with default parameters. Finally, the consensus sequence was polished once more using Hapo-G and short reads, to ensure correction of duplicate regions that were collapsed (Table S4).

### Validation of the *Triticum aestivum* cv Renan assembly

The quality value (QV) of the Renan and CS assemblies was obtained using Merqury[43]. First, 31-mers were extracted from the Renan and CS Illumina sequencing reads (accessions SRR5893651, SRR5893652, SRR5893653 and SRR5893654) and then the QV of each genome assembly was computed using Merqury (version 1.3, git commit 6b5405e). We used BLAST[25] to search for the presence of 107,891 HC genes from CS RefSeq v1.1 in the Renan genome sequence. We extracted the 461,476 individual exons larger than 30 bps and without Ns from this dataset and computed exon-by-exon BLAST in order to avoid spurious sliced alignments. An exon was considered present if it matched the Renan scaffolds with at least 95% identity over at least 95% of its length. To estimate the proportion of identical exons between CS and Renan and the average nucleotide identity, we used the same BLAST-based procedure but while restricting the dataset to 454,008 CS exons that are on pseudomolecules (excluding chrUn) and considering Renan pseudomolecules instead of scaffolds i.e., only exons carried by the same chromosome in CS and Renan were considered. We extracted all available ISBPs (150 bps each) from the CS RefSeq v1.1 and filtered out ISBPs containing Ns and those that do not map uniquely on the CS genome. This led to the design of a dataset containing 5,394,172 ISBPs which were aligned on the Renan scaffolds using BLAST. We considered an ISBP was conserved in Renan if it matched with at least 90% identity over 90% of its length. We used the same ISBP dataset to study the impact of polishing on error rate in the assembly while using BLAST and considering at least 90% identity over at least 145 aligned nucleotides.

### Anchoring of the *Triticum aestivum* cv Renan assembly

We guided the construction of 21 Renan pseudomolecules based on collinearity with the CS RefSeq Assembly v2.1. For this, we used the positions of conserved ISBPs as anchors (5,087,711 ISBPs matching with >=80% identity over >=90% query overlap). This represented 357 ISBPs/Mb, meaning that even the smallest scaffolds (30kb) carried generally more than 10 potential anchors. However, some ISBPs match at non-orthologous positions which create noise to precisely determine the order and orientation of some scaffolds. To overcome this issue, we considered ISBPs by pairs. Only pairs of adjacent ISBPs (i.e. separated by less than 50kb on both CS and Renan genomes) were kept as valid anchors, allowing the filtering out of isolated mis-mapped ISBPs. Only scaffolds harboring at least 50% of valid ISBP pairs on a single chromosome were kept. The others were considered unanchored and they comprised the “chrUn”. We calculated the median position of matching ISBP pairs along each CS chromosome for defining the order of the Renan scaffolds relative to each other. Their orientation was retrieved from the orientation of all matching ISBP pairs in CS following the majority rule. We thus built 21 pseudomolecules that were then corrected according to the HiC map as explained hereafter.

Two Hi-C biological replicates were prepared from ten-days plantlets of *Triticum aestivum* cv. Renan following the Arima Hi-C protocol (Arima Hi-C User Guide for Plant Tissues DOC A160106 v01). For each replicate, two libraries were constructed using the Kapa Hyper Prep kit (Roche) according to Arima’s recommendation (Library Preparation using KAPA Hyper Prep Kit DOC A160108 v01). The technical replicates were then pooled and sent to Genewiz for sequencing on an Illumina HiSeq4000 (four lanes in total), reaching a 35x coverage. We mapped a sample of 240 million read pairs with BWA-MEM (Burrows-Wheeler Aligner, Heng Li, 2013) to the formerly built 21 pseudomolecules, filtered out for low quality, sorted, and deduplicated using the Juicer pipeline[44]. We produced a Hi-C map from the Juicer output by the candidate assembly visualizer mode of 3D-DNA pipeline[45] and visualized it with the Juicebox Assembly Tools software. Based on abnormal frequency contacts signals revealing a lack of contiguity, scaffold-level modifications of order, orientation and/or chimeric scaffolds were identified in order to improve the assembly. In case of chimeric scaffolds, coordinates of resulting fragments were retrieved from the Juicebox Assembly Tools application but then recalculated to correspond precisely to the closest gap in the scaffold. Pseudomolecules were eventually rebuilt from initial scaffolds and new fragments while adding 100N gaps between neighbor scaffolds. A final Hi-C map was built to validate the accuracy of the final assembly.

### Calculation of chromosome coverage

Short (*Triticum aestivum* cv Renan and *Ae. ventricosa*) and long-reads (*Triticum aestivum* cv Renan) were aligned using minimap2 (with the following parameters ‘-I 17G −2 --sam-hit-only -a -x sr’ and ‘-I 17G −2 --sam-hit-only --secondary=no -a -x map-ont’ respectively). Coverage of individual chromosomes was calculated in 1 Mb windows using mosdepth[46] (version 0.3.1) and the following parameters ‘--by 1000000 -n -i 2 -Q 10 -m’. Note that the ‘-i 2’ and ‘-Q 10’ parameters were used to keep only alignments of reads that mapped in a proper pair and with a minimal quality value of 10. Coverage of individual chromosomes was plotted in Figure 1. In addition, large deletions and duplications were detected using CNVnator[47] with the Illumina bam file and a window of 100bp. We focused on large events (>500kb) and detected only 15 deletions and no duplication (Figure 1).

### Transposable elements annotation

Transposable elements were annotated using CLARITE[28]. Briefly, TEs were identified through a similarity search approach based on the ClariTeRep curated databank of repeated elements using RepeatMasker (www.repeatmasker.org) and modelled with the CLARITE program that was developed to resolve overlapping predictions, merge adjacent fragments into a single element when necessary, and identify patterns of nested insertions[28].

### Gene prediction

We used MAGATT pipeline (Marker Assisted Gene Annotation Transfer for Triticeae, https://forgemia.inra.fr/umr-gdec/magatt) to map the full set of 106,801 High Confidence and 159,848 Low Confidence genes predicted in Chinese Spring IWGSC RefSeq v2.1. The workflow implemented in this pipeline was described in Zhu et al.[22]. Briefly, it uses gene flanking ISBP markers in order to determine an interval that is predicted to contain the gene before homology-based annotation transfer, limiting problems due to multiple mapping. When the interval is identified, MAGATT uses BLAT[48] to align the gene (UTRs, exons, and introns) sequence and recalculate all sub-features coordinates if the alignment is full-length and without indels. If the alignment is partial or contains indels, it runs GMAP[49] to perform spliced alignment of the candidate CDS inside the interval. If no ISBP-flanked interval was determined or if both BLAT and GMAP failed to transfer the gene, MAGATT runs GMAP against the whole genome, including the unanchored fraction of the Renan assembly. We kept the best hit considering a minimum identity of 70% and a minimum coverage of 70%, with *cross_species* parameter enabled.

We then masked the genome sequence based on mapped genes and predicted transposable elements coordinates using BEDTools[50] mergeBed and maskfasta v2.27.1. Hence, we computed a *de novo* gene prediction on the unannotated part of the genome. We used TriAnnot[30] to call genes based on a combination of evidence: RNA-Seq data, *de novo* predictions of gene finders (FGeneSH, Augustus), similarity with known proteins in *Poaceae*, as described previously[7]. For that purpose, we mapped RNA-Seq reads with hisat2[51] v2.0.5, called 277,505 transcripts with StringTie[52] v2.0.3, extracted their sequences with Cufflink[53] gffread v2.2.1, and provided this resource as input to TriAnnot. We optimized TriAnnot workflow to ensure a flawless use on a cloud-based hpc cluster (10 nodes with 32 CPUs/128GB RAM each and shared file system) using the IaaS Openstack infrastructure from the UCA Mesocentre. Gene models were then filtered as follows: we discarded gene models that shared strong identity (>=92% identity, >=95% query coverage) with an unannotated region of the Chinese Spring RefSeq v2.1, considered as doubtful predictions. We then kept all predictions that matched RNASeq-derived transcripts (>=99% identity, >=70% query and subject coverage). For those that did not show evidence of transcription, we kept gene models sharing protein similarity (>=40% identity, >=50% query and subject coverage) with a *Poaceae* protein having a putative function (filtering out based on terms “unknown”, “uncharacterized”, and “predicted protein”).

### Comparison of genome assemblies

Genome assemblies were downloaded from https://webblast.ipk-gatersleben.de/downloads. Contigs were extracted by splitting input sequences at each N and standard metrics were computed. Gene completion metrics were calculated using BUSCO v5.0 and version 10 of the poales geneset which contains 4896 genes.

We built dotplots between Renan, CS and 10 other reference quality genomes (Arina*LrFor*, CDC Landmark, CDC Stanley, Jagger, Julius, LongReach Lancer, Mace, Norin61, SY Mattis, spelta PI190962) by using orthologous positions of conserved ISBPs (1 ISBP every 2.5kb on average) identified by mapping them with BWA-MEM (maximum 2 mismatches, 100% coverage and minimal mapping quality of 30).

### Characterisation of haplotypic blocks

First a colored de Bruijn graph was built for each chromosome from the eleven available chromosome-scale assemblies of wheat (Renan, CS, Arina*LrFor*, CDC Landmark, CDC Stanley, Jagger, Julius, LongReach Lancer, Mace, Norin61 and SY Mattis). The colored de Bruijn graph was created using Bifrost[54] with 31-mers and a unique color for each wheat cultivar. In a second step, we extracted short markers (1kb) evenly spaced (20kb or 5kb) on each chromosome and queried the colored de Bruijn graph using Bifrost and the following parameter ‘-e 0.95’ (for the comparison of each chromosome) and ‘-e 0.97’ (for the comparison of the *Sm1* locus). This parameter is the ratio of k-mers from queries that must occur in the graph to be reported as present. For whole chromosome analyses, the 20kb blocks were merged into 1-Mb blocks (the most abundant colour in the 50 20kb blocks was retained for the 1Mb block). Individual blocks and *Ae. ventricosa* coverage were displayed using RIdeogram[55].

### Comparison of a storage protein coding gene cluster

We performed manual curation of the gene models encoding storage proteins predicted in Renan. Protein sequences of prolamin and resistance genes[35] from a 1B chromosome locus were downloaded and aligned to the CS and Renan genomes using BLAT[48] with default parameters. Draft alignments were refined by aligning the given protein sequence and the genomic region defined by the blat alignment using Genewise with default parameters. Resulting alignments were filtered in order to conserve only the best match for each position by keeping only the highest-scoring alignment and the genomic region containing the gene cluster was extracted. Then, we used the jcvi suite[56] with the mcscan pipeline to find synteny blocks between both genomes. First, we used the “jcvi.compara.catalog” command to find orthologs and then the “jcvi.compara.synteny mcscan” with “--iter=1” command to extract synteny blocks. Finally, we generated the figure with the “jcvi.graphics.synteny” command and manually edited the generated svg file to improve the quality of the resulting image by changing gene colors, incorporating gaps and renaming genes. Moreover, to make the figure clearer, we artificially reduced the intergenic space by 95% so that gene structures appear bigger. The omega gene cluster representation figure was generated by using DnaFeaturesViewer[57] with coordinates of features generated by the mcscan pipeline used previously.

## Additional files

All the supporting data are included in two additional files: (a) A supplementary file which contains Supplementary Tables 1-7 and Supplementary Figures 1-3; (b) A supplementary file which contains dotplots of the 21 chromosomes of Renan with other wheat genome assemblies.

## Acknowledgements

This work was supported by the Genoscope, the Commissariat à l’Énergie Atomique et aux Énergies Alternatives (CEA) and France Génomique (ANR-10-INBS-09-08). The biological material (i.e. plant production, sample management, DNA and RNA extractions performed by Caroline Pont and Cécile Huneau at GDEC) have been obtained in the framework of the France Génomique WheatOMICS project (2017-2021) coordinated by Jérôme Salse. The authors are grateful to Oxford Nanopore Technologies Ltd for providing early access to the PromethION device through the PEAP, and we thank the staff of Oxford Nanopore Technologies Ltd for technical help. We are grateful to the Mésocentre Clermont Auvergne University and/or AuBi platform for providing help and/or computing and/or storage resources.

## Availability of supporting data

The Illumina and PromethION sequencing data and the Bionano optical map are available in the European Nucleotide Archive under the following project PRJEB49351. The genome assembly and gene predictions are freely available from the Genoscope website http://www.genoscope.cns.fr/plants/.

Additionally, all the data and scripts used to produce the main figures are available on a github repository https://github.com/institut-de-genomique/Renan-associated-data

## Competing interests

The authors declare that they have no competing interests. JMA received travel and accommodation expenses to speak at Oxford Nanopore Technologies conferences. JMA and CB received accommodation expenses to speak at Bionano Genomics user meetings.

## Funding

This work was supported by the Genoscope, the Commissariat à l’Énergie Atomique et aux Énergies Alternatives (CEA) and France Génomique (ANR-10-INBS-09-08). CM postdoc was funded by the SRESRI/Région Auvergne-Rhône-Alpes.

## Author’s contributions

SA, ID and AB extracted the sequenced DNA and generated the optical map. KL and AA optimized and performed the nanopore and Illumina sequencing. NP, EP and MR generated the Hi-C libraries and sequences. JMA, SE, BI, CM, PLZ, CB, HR, PL, DG and FC performed the bioinformatic analyses. JMA, SE, BI, CM, PLZ, CB, CC, HR, PL and FC wrote the article. JMA, PW and FC supervised the study.

## Notes

http://www.genoscope.cns.fr/plants/

### Competing Interest Statement

The authors declare that they have no competing interests. Jean-Marc Aury received travel and accommodation expenses to speak at Oxford Nanopore Technologies conferences. Jean-Marc Aury and Caroline Belser received accommodation expenses to speak at Bionano Genomics user meetings.

