## Supplementary Data for "Long-read and chromosome-scale assembly of the hexaploid wheat genome achieves high resolution for research and breeding"

**Table S1: Statistics of the ONT sequencing data**

| | Raw ONT reads | Longest reads |
|---|---|---|
| Number of runs | 2 MinION + 18 PromethION | |
| Cumulative size | 1,074 Gb | 510 Gb |
| # of reads | 120,883,721 | 11,654,667 |
| Coverage (17Gb) | 63 X | 30 X |
| N50 (bp) | 24,651 | 45,077 |
| Coverage (>50Kb) | 14.1 X | 14.1 X |

**Table S2: BNG Optical map**

| | DLE-1 genome map |
|---|---:|
| Genome map number | 1,053 |
| Total Genome Map Length (Mbp) | 14,947 |
| Genome Map N50 (Mbp) | 37.5 |

**Table S3: Raw long-read assemblies**

| | SMARTDeNovo | Redbean | Flye |
|---|---|---|---|
| Subset of reads used | Longest - 30X | Longest - 30X | Longest - 30X |
| Cumulative size | **14,155,561,280** | 29,695,324,870 | 13,050,586,822 |
| # contigs | 21,197 | 646,779 | **14,205** |
| longest contig (bp) | 13,367,134 | 957,564 | **18,693,219** |
| minimal size (bp) | 34,696 | 1,157 | 50,033 |
| N50 (L50) | 1,142,720 (3,758) | 76,231 (107,967) | **1,775,593 (2,165)** |
| N90 (L90) | 322,974 (12,488) | 21,181 (397,125) | **471,502 (7,566)** |
| Restitution time (h) | 408 | 168 | 1,032 |
| IT | 8 nodes with 32-core and 1.5Tb | 1 node of 64-core and 3Tb | 1 node of 64-core and 3Tb |

**Table S4: Hybrid assemblies obtained using ONT and BNG data**

| | | SMARTDeNovo | Flye |
|---|---|---|---|
| Scaffolds | Cumulative size | **14,272,571,061** | 14,204,637,528 |
| | # scaffolds | 4,255 | **4,135** |
| | longest scaffold (bp) | **253,607,211** | 252,437,931 |
| | N50 (L50) | 48,460,876 (79) | **49,688,204 (77)** |
| | N90 (L90) | 7,877,958 (378) | **8,121,250 (362)** |
| | % of unknown bases | **1.81%** | 5.99% |
| Contigs | Cumulative size | **14,014,403,266** | 13,330,835,323 |
| | # contigs | **14,241** | 15,282 |
| | longest contig (bp) | 15,116,800 | **19,215,531** |
| | N50 (L50) | **2,144,699 (1,963)** | 1,662,574 (2,131) |
| | N90 (L90) | **595,732 (6,672)** | 440,639 (8,204) |
| | BUSCO<br>(n=4,896) | **Complete : 97.0%**<br>**Duplicated : 87.3%**<br>**Fragmented : 0.4%**<br>**Missing : 2.6%** | Complete : 96.7%<br>Duplicated : 86.6%<br>**Fragmented : 0.4%**<br>Missing : 2.9% |

**Table S5: Impact of the polishing**

| | | SMARTDeNovo raw | SMARTDeNovo polished with long reads | SMARTDeNovo polished with short reads |
|---|---|---|---|---|
| BUSCO (n=4,896) | Complete | 83.0% | 96.7% | 96.6% |
| | Duplicated | 32.5% | 83.1% | 87.0% |
| | Fragmented | 2.4% | 0.6% | 0.6% |
| | Missing | 14.6% | 2.7% | 2.8% |
| IBSPs (n=5.76 M) | Aligned | 80.4% | 92.9% | 93.4% |
| | Perfectly aligned | 7.0% | 28.0% | 58.9% |

**Table S6: Statistics of the ONT reads obtained with two different versions of the guppy basecaller**

| | Guppy 3.3 | Guppy 3.6 |
|---|---|---|
| Number of runs | 2 MinION + 18 PromethION<br>Selection of the longest reads (30X) | |
| Cumulative size | 510 Gb | 510 Gb |
| # of reads | 11,731,678 | 11,654,667 |
| Coverage (17Gb) | 30 X | 30 X |
| N50<br>(bp) | 44,797 | 45,077 |
| Coverage (>50Kb) | 13.9 X | 14.1 X |

**Table S7: Long-read assemblies with ONT reads obtained with two different versions of the guppy basecaller**

| | SMARTDeNovo | SMARTDeNovo |
|---|---|---|
| Subset of reads used | Guppy 3.3<br>Longest - 30X | Guppy 3.6<br>Longest - 30X |
| Cumulative size | 13,975,909,499 | **14,155,561,280** |
| # contigs | 26,167 | **21,197** |
| longest contig (bp) | 6,593,358 | **13,367,134** |
| minimal size (bp) | 33,360 | **34,696** |
| N50 (L50) | 899,898 (4,685) | **1,142,720 (3,758)** |
| N90 (L90) | 247,425 (15,801) | **322,974 (12,488)** |

**Table S8: RNASeq data for Renan from 28 samples corresponding to 14 different organs/conditions in replicates.**

| Samples accession | Genotype | Zadoks condition | Type | Tissue | Wheat Ontology (CO) | Plant Ontology (PO) | Stress Ontology (PSO) |
|---|---|---|---|---|---|---|---|
| SAMEA11349165 | Renan | 100DD | control | Grain | CO_321:0001679 | PO:0030104 | |
| SAMEA11349166 | Renan | 500DD | control | Grain | CO_321:0001679 | PO:0030104 | |
| SAMEA11349167 | Renan | 250DD | control | Grain | CO_321:0001679 | PO:0030104 | |
| SAMEA11349168 | Renan | 500DD | control | Grain | CO_321:0001679 | PO:0030104 | |
| SAMEA11349169 | Renan | 250DD | control | Grain | CO_321:0001679 | PO:0030104 | |
| SAMEA11349170 | Renan | 100DD | stress | Grain | CO_321:0001679 | PO:0030104 | PSO:0000012 |
| SAMEA11349171 | Renan | 500DD | stress | Grain | CO_321:0001679 | PO:0030104 | PSO:0000012 |
| SAMEA11349172 | Renan | 100DD | stress | Grain | CO_321:0001679 | PO:0030104 | PSO:0000012 |
| SAMEA11349173 | Renan | 100DD | control | Grain | CO_321:0001679 | PO:0030104 | |
| SAMEA11349174 | Renan | 250DD | stress | Grain | CO_321:0001679 | PO:0030104 | PSO:0000012 |
| SAMEA11349175 | Renan | 500DD | stress | Grain | CO_321:0001679 | PO:0030104 | PSO:0000012 |
| SAMEA11349176 | Renan | 250DD | stress | Grain | CO_321:0001679 | PO:0030104 | PSO:0000012 |
| SAMEA11349177 | Renan | 700DD | control | Grain | CO_321:0001679 | PO:0030104 | |
| SAMEA11349178 | Renan | 700DD | control | Grain | CO_321:0001679 | PO:0030104 | |
| SAMEA11349179 | Renan | 700DD | stress | Grain | CO_321:0001679 | PO:0030104 | PSO:0000012 |
| SAMEA11349180 | Renan | 700DD | stress | Grain | CO_321:0001679 | PO:0030104 | PSO:0000012 |
| SAMEA11349181 | Renan | Z13 | control | Root | CO_321:0000476/10 | PO:0009005 | |
| SAMEA11349182 | Renan | Z13 | control | Root | CO_321:0000476/10 | PO:0009005 | |
| SAMEA11349183 | Renan | Z13 | control | Leaves | CO_321:0000476/29 | PO:0025034 | |
| SAMEA11349184 | Renan | Z13 | control | Leaves | CO_321:0000476/29 | PO:0025034 | |
| SAMEA11349185 | Renan | Z32 | control | Leaves | CO_321:0000476/29 | PO:0025034 | |
| SAMEA11349186 | Renan | Z32 | control | Leaves | CO_321:0000476/29 | PO:0025034 | |
| SAMEA11349187 | Renan | Z61 | control | Leaves | CO_321:0000476/29 | PO:0025034 | |
| SAMEA11349188 | Renan | Z61 | control | Leaves | CO_321:0000476/29 | PO:002503 | |
| SAMEA11349189 | Renan | Z61 | control | Stem | CO_321:0000476/46 | PO:0009047 | |
| SAMEA11349190 | Renan | Z61 | control | Stem | CO_321:0000476/46 | PO:0009047 | |
| SAMEA11349191 | Renan | Z32 | control | Stem | CO_321:0000476/46 | PO:0009047 | |
| SAMEA11349192 | Renan | Z61 | control | Stem | CO_321:0000476/46 | PO:0009047 | |

**Figure S1.** Validation of both ends, which contains telomeric repeats, of the chromosome 7A The optical maps are in blue and the chromosome sequence in green. Restriction sites are represented by vertical lines and are joined between the sequence and the map when properly aligned..

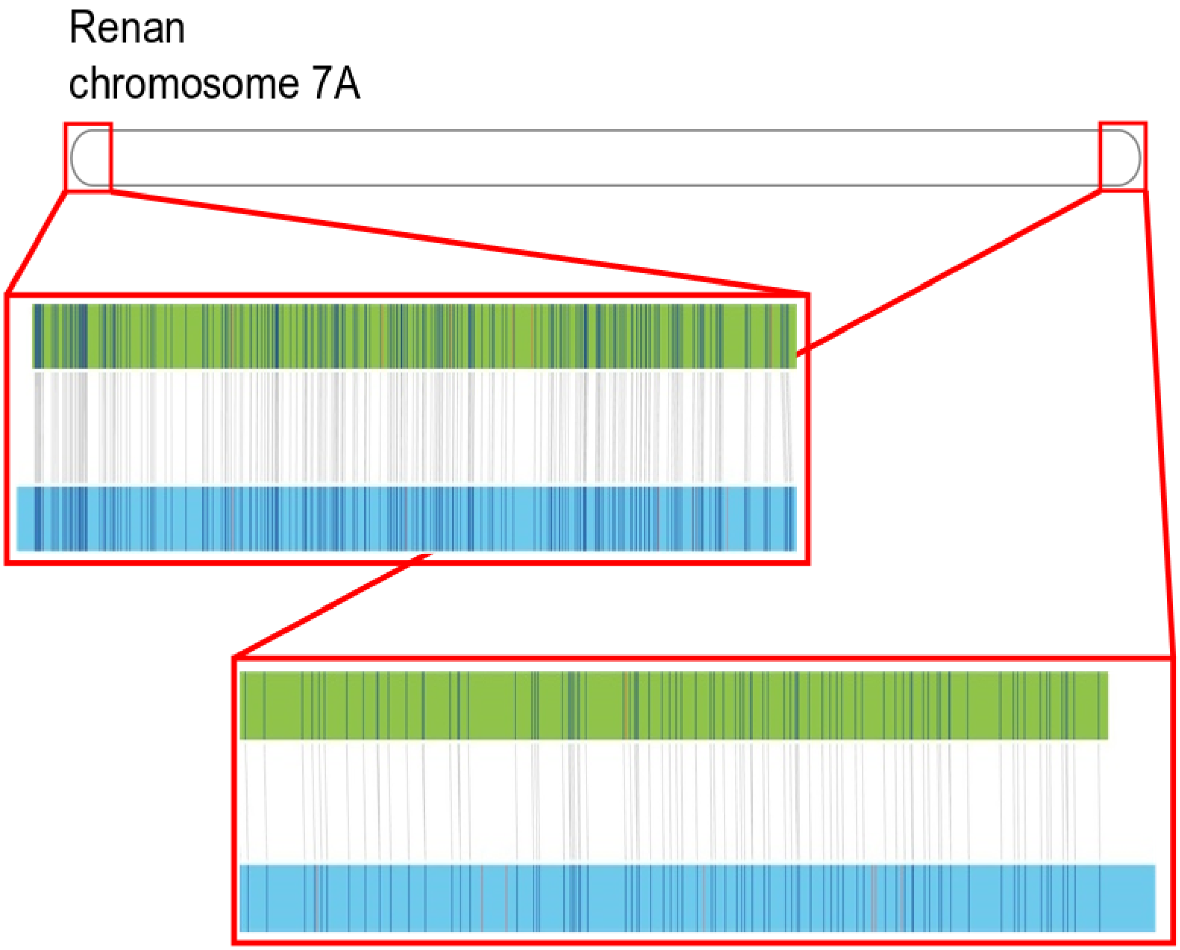

**Figure S2.** Comparison of the accuracy of different ONT basecallers. **A.** ONT reads from a yeast sample. **B.** ONT ultra-long reads (>100 kb) from a wheat sample.

**A.**

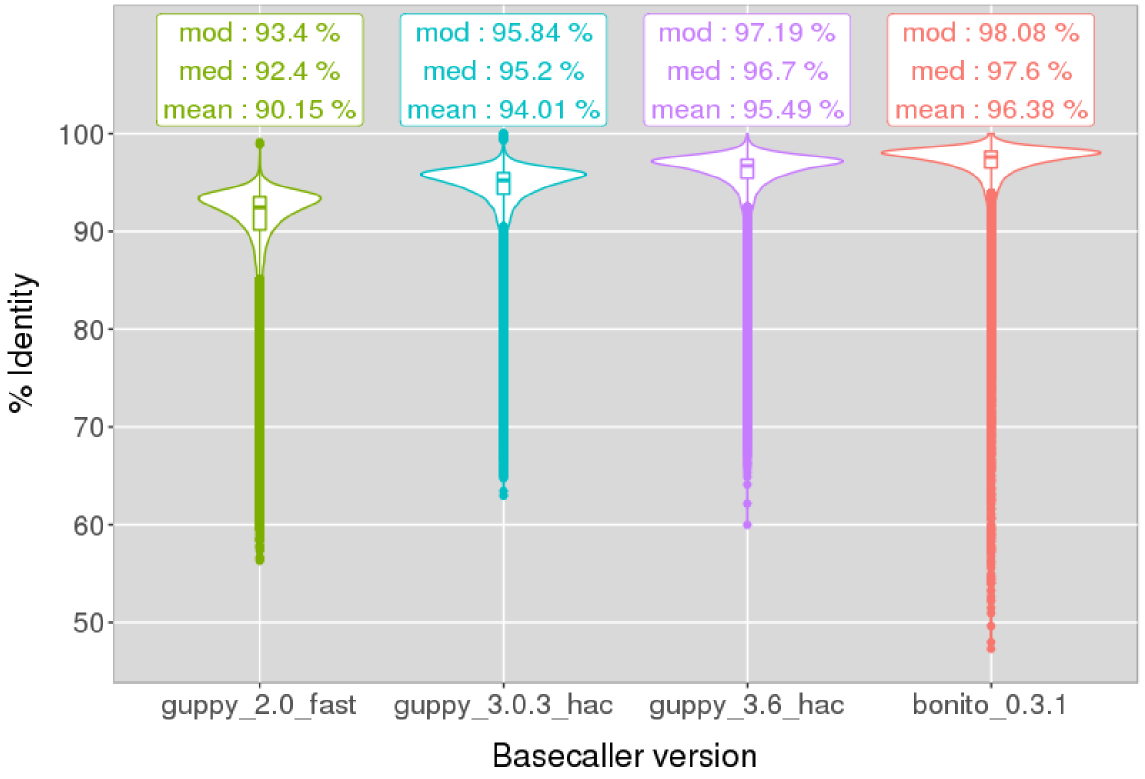


**B.**

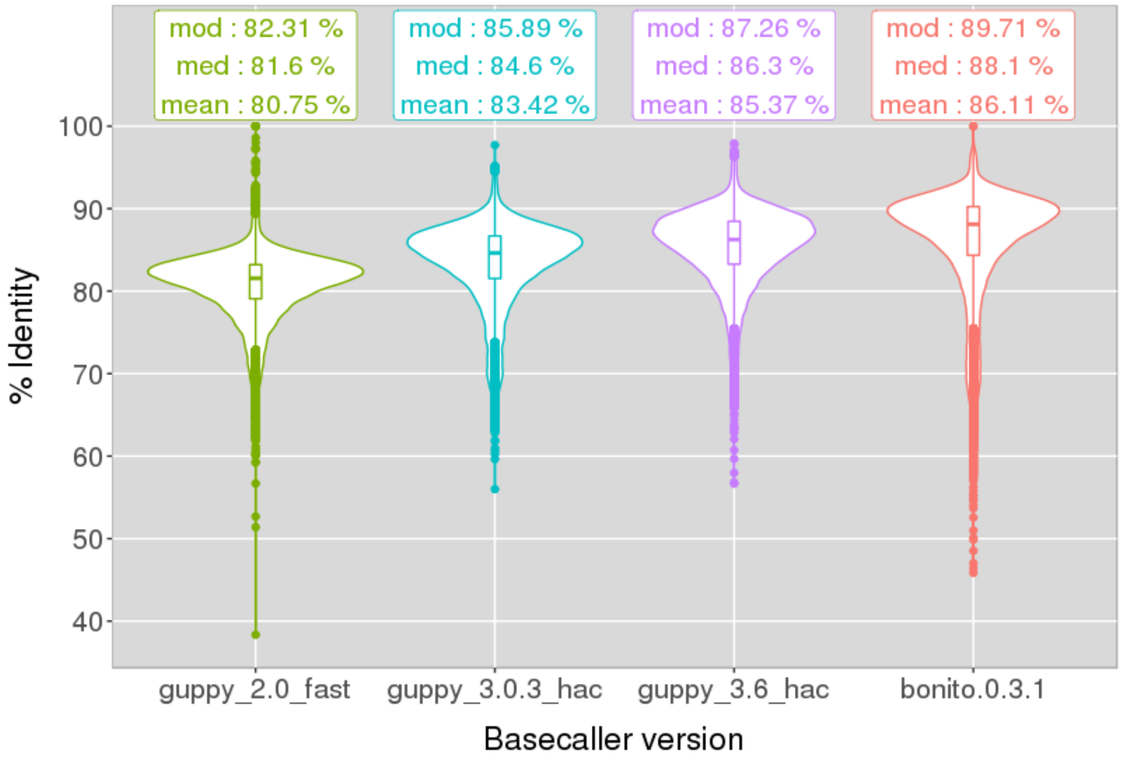

**Figure S3.** Number of gaps per Mbp in Chinese Spring and Renan genome assemblies.

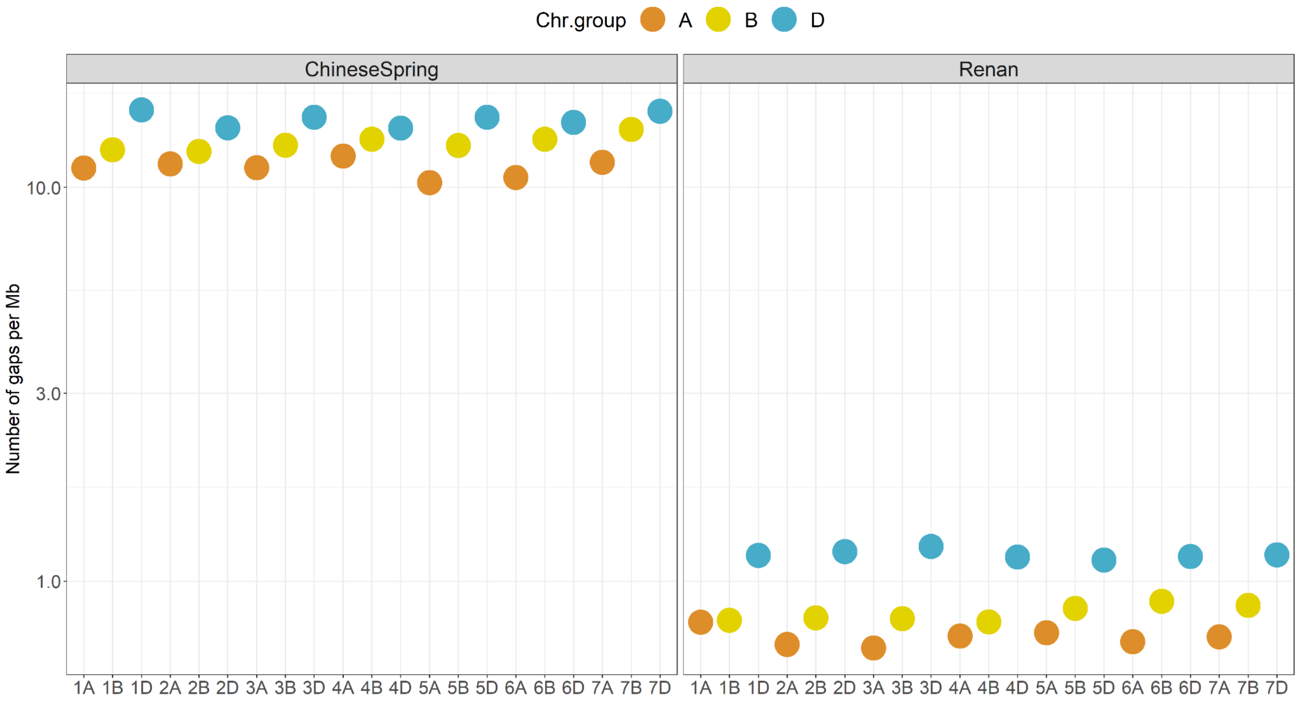

**Figure S4.** Large-scale structural variation in chromosomes 5B and 7B of ArinaLrFor and SY Mattis cultivars. Translocated regions correspond to the blocks in cyan (chromosome 5B) and green (chromosome 7B) that appear specific to SY Mattis and ArinaLrFor.

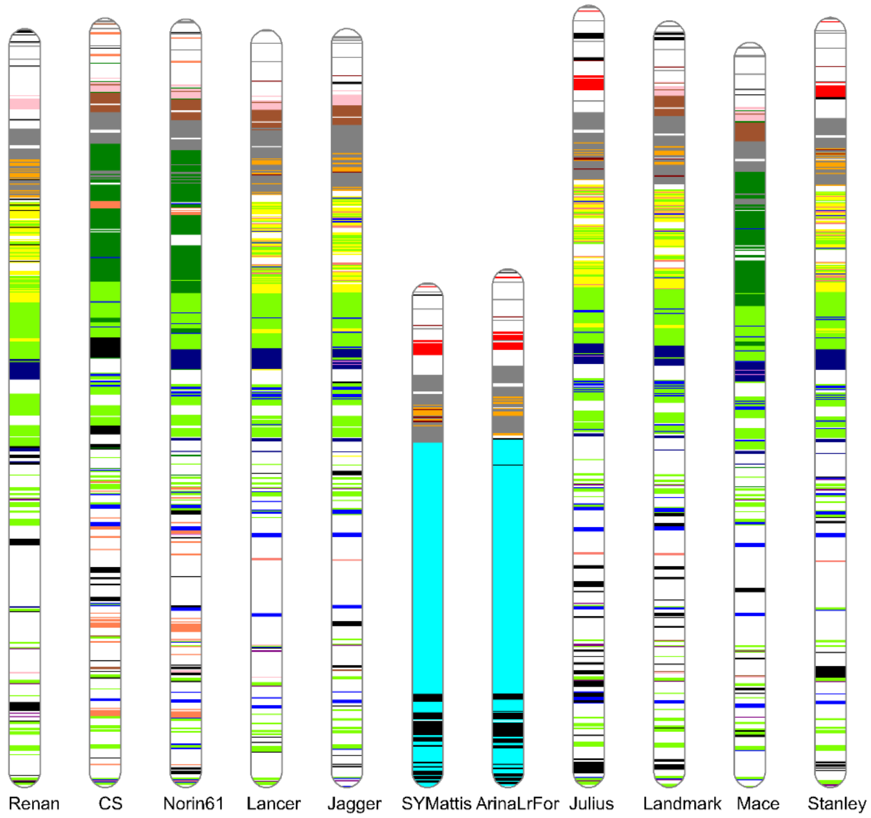


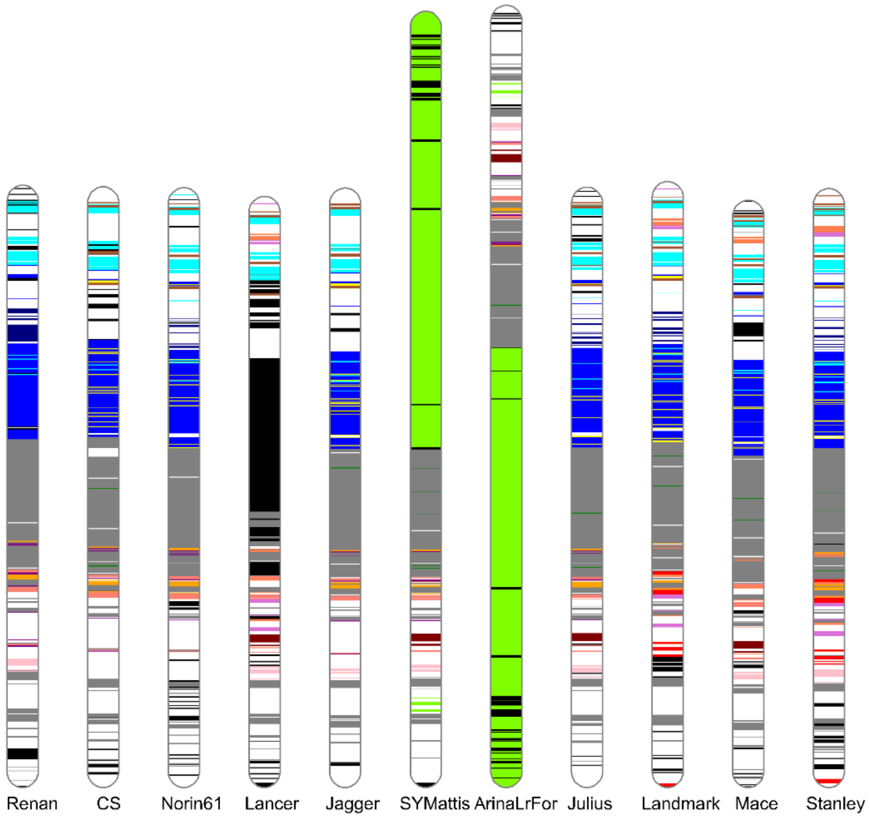
